# MillionTrees: A global benchmark for tree detection across geography, sensor, and annotation geometry

**DOI:** 10.64898/2026.10.30.762845

**Authors:** Ben G. Weinstein, Josh Veitch-Michaelis, Matt J. Allen, Adison Altamirano, Sarah Anacleto, Rasmus Astrup, James G. C. Ball, Hugo Baudchon, Sara Beery, Stephanie A. Bohlman, John Brandt, Myriam Cloutier, Simon-Olivier Duguay, Jessica Ertel, Zhengpeng Feng, Adnan Firoze, Julian Frey, G. Andrew Fricker, Dimitri Gominski, Sarah Graves, Stuart W. D. Grieve, Sander de Haas, Sebastian H. M. Hickman, Tobias D. Jackson, Andrew Jansen, Teja Kattenborn, Taimur Khan, Ryoungseob Kwon, Kyaw Kyaw Htoo, Etienne Laliberté, Isabelle Lefebvre, Emily R. Lines, Moritz Lucas, Alejandro Miranda, Blas Mola-Yudego, Maurice Mugabowindekwe, Jaylen Nicholson, Masanori Onishi, Yusuke Onoda, Arthur Ouaknine, Harry J. F. Owen, Camille Pawlak, Stefano Puliti, Md. Farhadur Rahman, David J. Russell, Oliver Schütte, Henry Senyondo, Yihang She, Ying Sun, Ryuichi Takeshige, Jonas Troles, Vicente Vasquez, Jonathan Ventura, Benjamin Wagner, Qinchuan Xin, Qiue Xu, Derek J. N. Young, Pedro Zamboni, Ethan P. White

**Affiliations:** Department of Wildlife Ecology and Conservation, University of Florida, Gainesville, FL, USA; Department of Geography, University of Cambridge, Cambridge, UK; Laboratorio de Ecología del Paisaje y Conservación, Universidad de La Frontera, Temuco, Chile; Norwegian Institute for Bioeconomy Research (NIBIO), Division of Forest and Forest Resources, Høgskoleveien 8, 1433 Ås, Norway; Conservation Research Institute, Department of Plant Sciences, University of Cambridge, Cambridge, UK; School of Biological Sciences, University of Bristol, 24 Tyndall Avenue, Bristol BS8 1TQ, UK; Institut de recherche en biologie végétale, Université de Montréal, 4101 Sherbrooke Est, Montréal, Canada; Mila - Québec AI Institute, 6666 St-Urbain, Montréal, Canada; Office of the Chief Remote Pilot, Parks Australia, Department of Climate Change, the Environment and Energy, Parkes ACT 2600, Australia; Massachusetts Institute of Technology, Cambridge, MA, USA; World Resources Institute, 10 G St NE #800, Washington, DC 20002, USA; Département de sciences biologiques, Université de Montréal, Montréal, QC, Canada; Department of Computer Science and Technology, University of Cambridge, Cambridge, UK; Department of Computer Science, Purdue University, West Lafayette, IN, USA; Department of Social Sciences, California Polytechnic State University, San Luis Obispo, CA, USA; Department of Geosciences and Natural Resource Management, University of Copenhagen, Copenhagen, Denmark; Department of Geography and Environmental Science, Queen Mary University of London, UK; Digital Environment Research Institute, Queen Mary University of London, UK; Justdiggit, 1012 TM Amsterdam, The Netherlands; Yusuf Hamied Department of Chemistry, University of Cambridge, Lensfield Road, Cambridge, CB2 1EW, UK; Helmholtz Centre for Environmental Research - UFZ, Theodor-Lieser-Str. 4, 06120 Halle (Saale), Germany; Seoul National University, Seoul, South Korea; Graduate School of Agriculture, Kyoto University, Kyoto, Japan; Field Science Center for Northern Biosphere, Hokkaido University, Hokkaido, Japan; Joint Lab Artificial Intelligence and Data Science, Osnabrück University, Osnabrück, Germany; Laboratorio de Observación de la Tierra y Ecología de Bosques, Departamento de Ciencias Forestales, Universidad de La Frontera, Temuco, Chile; Landscape and Carbon Dynamics CoLab, School of Forest Sciences, University of Eastern Finland, Joensuu, Finland; Research School of Biology, Australian National University, Canberra ACT 0200, Australia; DeepForest Technologies Co., Ltd., Kyoto, Japan; McGill University, Montréal, Québec, Canada; Department of Geography, University of California, Los Angeles, Los Angeles, CA, USA; Norwegian Institute for Bioeconomy Research (NIBIO), Høgskoleveien 7, 1433 Ås, Norway; Chair of Sensor-based Geoinformatics (geosense), University of Freiburg, Germany; Chair of Forest Growth and Dendroecology, University of Freiburg, Germany; Department of Remote Sensing and GIS, Gazipur Agricultural University, Dhaka, Bangladesh; Department of Plant Sciences, University of California, Davis, Davis, CA, USA; Working Group Remote Sensing and Digital Image Analysis, Osnabrück University, Osnabrück, Germany; School of Geography and Planning, Sun Yat-sen University, Guangzhou, China; Graduate School of Science, Osaka Metropolitan University, Osaka, Japan; Cognitive Systems Group, University of Bamberg, Bamberg, Germany; Department of Computer Science & Software Engineering, California Polytechnic State University, San Luis Obispo, CA, USA; Huanjiang Observation and Research Station for Karst Ecosystem, Guangxi Key Laboratory of Karst Ecological Processes and Services, Institute of Subtropical Agriculture, Chinese Academy of Sciences, Changsha 410125, China; University of Chinese Academy of Sciences, Beijing 100049, China; The University of Melbourne, 500 Yarra Boulevard, Richmond, VIC 3121, Australia; Dresden University of Technology, Dresden, Germany; Department of Earth Observation, Rwanda Space Agency, Kigali, Rwanda; School of Forest, Fisheries and Geomatics Sciences, University of Florida, Gainesville, FL, USA; Department of Forest and Wildlife Ecology, University of Wisconsin-Madison, Madison, WI, USA; Aspia Space, FibreHub R2, Trevensen Lane, Pool, Cornwall, TR15 3GF, UK

## Abstract

Airborne tree monitoring using remotely sensed data, combined with the growth of computer vision models for object detection and segmentation, can facilitate broad scale assessment of tree health, ecology and ecosystem function. However, datasets for airborne tree detection have been largely limited to a single geography, resolution, or annotation approach, making it difficult to develop general models and benchmark their utility for broad applications. We compile existing and newly published annotation datasets to create a comprehensive dataset for training and evaluating airborne tree detection in RGB imagery. The MillionTrees dataset has over a million individual trees from a broad range of forest types, tree species, and geographic locations. In addition, it has six million weakly supervised annotations from a range of sources including airborne LiDAR and photogrammetry, arborist surveys, and simulated landscapes. An evaluation standard is provided to facilitate rapid model development. We benchmark several existing tree detection algorithms and demonstrate that while current open-source models do not generalize across the diverse datasets in the benchmark, training on MillionTrees improves both within-domain and out-of-domain performance. Held-out validation datasets using ground truth derived from terrestrial LiDAR paired with overhead RGB show that overfitting to test data is common and hyperparameter tuning is an underappreciated challenge towards a global zero shot tree detection method. We discuss the limitations of the currently available data for airborne RGB tree detection, how to create more robust evaluation data and the need for better understanding of the uncertainty and utility of tree detection from airborne remote sensing.

## Introduction

Individual trees are a fundamental unit for forest monitoring, silviculture, urban planning and quantification of ecosystem services. Surveys of individual trees over large spatial scales are increasingly applied using high resolution remote sensing from drones, planes, and high resolution satellites (Wang et al. 2020, Brandt et al. 2020, Weinstein et al. 2024). To convert sensor data into ecological information, computer vision models learn features for detecting trees and delineating their boundaries using annotated image data (Ball et al., 2023, Weinstein et al. 2019, Khan et al. 2025). This detection and delineation of trees is a core challenge in environmental remote sensing, with many models, reviews, datasets and applications to capture information about individual trees from airborne perspectives including tree counts (Amirkolaee et al., 2023), crown segmentations (Aubry-Kientz et al., 2021), phenology (Kleinsmann et al., 2023), foliar traits (Marconi et al. 2021), and carbon sequestration (Juan-Ovejero et al., 2023). The accuracy of both the models and ground truth in this area remain unclear due to a lack of field-verified data, confusion on nomenclature and annotation approaches, as well as limitations in resolving overlapping tree crowns in dense canopy and obscured subcanopy trees (Allen et al. 2025).

Despite the extensive work in airborne tree detection and multiple open source implementations (Ball et al., 2023; Weinstein et al., 2021, Baudchon et al. 2026, Khan et al. 2025), existing approaches may struggle with generalization to novel regions, forest types, sensors, and resolutions (Weinstein et al. 2020, Gan et al. 2023, Graves et al. 2023). While many models are developed for a specific use case, users often apply them outside of their initial domain. Challenges with generalization are due in part to insufficiently large, diverse, accurately annotated datasets for training and evaluating the generalized computer vision models (Kattenborn et al. 2022, Doerksen and Kerner 2026). These data limitations are compounded by the lack of a standardized strategy for labeling individual trees within sensor data, which can make it difficult to use compiled datasets and result in varying accuracy of the associated ground truth data (Ke and Quackenbush, 2011; Pulido et al., 2020, Allen et al. 2025, Khan et al. 2026). There is a need for datasets that include: 1) a diverse range of geographic areas, sensors, resolutions, and annotation types; 2) open data for both airborne sensors and annotations; and 3) has a standardized evaluation structure to facilitate comparisons among models and evaluation of how well they generalize to novel contexts.

Here we develop a large-scale tree detection benchmark to train and assess general models for airborne tree detection using a combination of field-verified, image-annotated, simulated and weakly labeled data. In order to capture the range of trees at a global scale, we combine data from many efforts that use a range of annotation strategies. The MillionTrees benchmark will allow users to address a number of unanswered questions:

1. Can current models generalize across sensor resolutions, illumination conditions and imaging platforms?
2. Does combining data from different efforts with varying annotation protocols lead to better generalization?
3. How can we combine multiple annotation geometries and incomplete annotations to strengthen tree detection models?
4. What types of data are needed to strengthen the validation rigor for airborne tree detection?

This is made possible by collating a large number of annotations from around the world into a single format. We combine data from previous efforts, including the Auto-arborist dataset (Beery et al., 2022), the Open-Aerial Map Tree Canopy Datasets (Veitch-Michaelis et al., 2024), the NeonTreeEvaluation dataset (Weinstein et al., 2021), ReForestTree (Reiersen et al., 2022), SelvaBox (Baudchon et al., 2026), and NIBIO’s UAV tree damage dataset (Puliti and Astrup 2025), add new datasets, previously unreleased datasets, and provide an evaluation structure that allows for easily assessing model performance on a range of tasks. All annotations, airborne data, and evaluation code are readily accessible for model development using an open-source Python package.

## Results

### Dataset

We collected over a million annotations for point, box and polygon geometries (Table 1, Figure 1). The standardized structure, central download location, and ease of evaluation along with semantic versioning, will allow rapid development, prototyping and model comparison of tree detection algorithms. In addition, the dataset includes over six million weak labels generated using LiDAR-derived tree predictions as well as labels from simulated forests to facilitate pretraining. Given the large size of the dataset, we also release a ‘mini’ version with a single image per source to develop workflows, and a ‘small’ version limited to 50 images per source for rapid iteration. We do not track the performance of mini or small version releases. We report evaluation metrics for a held-out test set used during model development, as well as a final validation dataset that is run a single time to reduce potential hyperparameter overfitting (Figure 2). The validation datasets come from paired terrestrial LiDAR and airborne RGB co-registered data used to generate gold standard ground truth annotations combining below canopy and above canopy views followed by human review.

**Figure 1.**
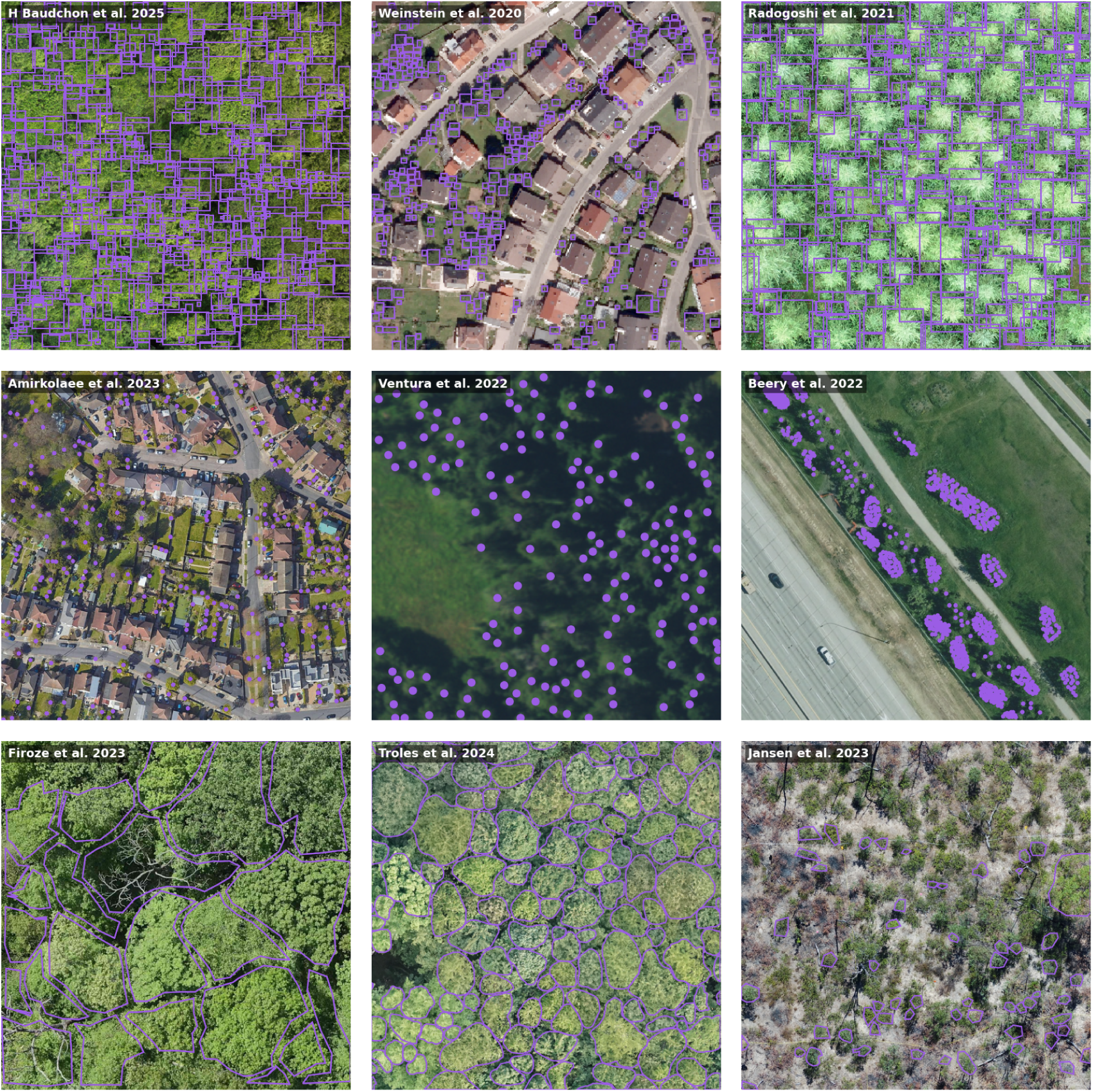
Sample images from the MillionTrees dataset. Point, box and polygon annotations for sources are overlaid on RGB airborne imagery.

**Figure 2.**
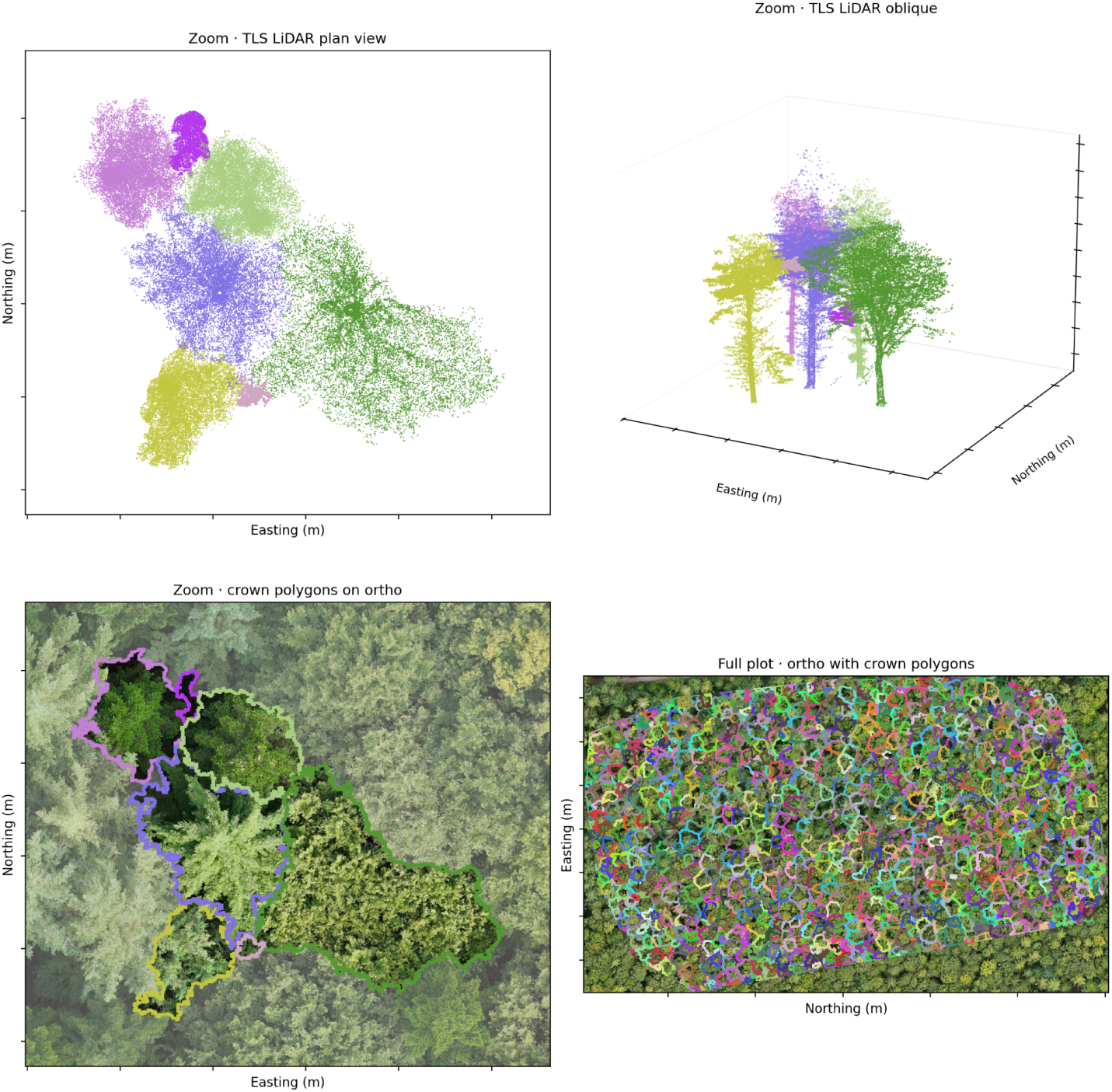
Validation dataset derived from terrestrial LIDAR scans paired with georegistered airborne RGB imagery from Frey et al. 2026. A sample set of annotations are shown to illustrate the connection between the point cloud, top down view and in the RGB orthomosaic. A supervised segmentation model Nguyen et al. 2026 was used to generate individual tree predictions in the point cloud, which was then reviewed by hand to align with the RGB data and correct oversegmentations that split a crown into multiple individuals.

**Table 1.** The number and diversity of source of annotation views for each dataset. The count of annotations differs from the individual ground truth trees in Table 6 due to multiple airborne surveys over the same area and edge overlap among source images that have been subdivided into smaller pieces.

| Dataset | Images | Annotations | Sources | Version |
|---|---|---|---|---|
| TreeBoxes | 11,968 | 945,022 | 12 | v0.23 |
| TreePoints | 35,103 | 916,030 | 10 | v0.23 |
| TreePolygons | 15,001 | 453,842 | 26 | v0.23 |
| **Subtotal** | 62,072 | 2,314,894 | 48 | v0.23 |
| Weak TreeBoxes | 45,717 | 6,413,695 | 1 | v0.23 |
| Weak TreePoints | 11,712 | 101,447 | 1 | v0.23 |
| Weak TreePolygons | 12,577 | 349,853 | 1 | v0.23 |

The MillionTrees dataset covers a range of backgrounds and geographies with a predominance of forested and urban landscapes (Figure 1). There is relatively little data from agricultural areas and transition zones with human habitation and dense trees. The ground resolution of imagery ranges from very high resolution drone imagery at less than 2 cm pixels to coarse imagery acquired by high elevation aircraft at 0.3m and above. The habitats and tree taxonomy generally favor temperate forests (e.g. Troles et al. 2024), but also include tropical forest examples (Baudchon et al. 2026, Vasquez et al. 2023). Arid areas are under-represented, but include a handful of datasets covering desert or scrub ecotypes (e.g. Jansen et al. 2023). While species labels are not included in the data release, data from over 20 countries and 6 continents ensures that the dataset provides a good foundation of variation in tree forms at a global scale.

### Benchmark model performance

Using existing models from the tree detection literature we assessed point, box, and polygon prediction accuracy for the MillionTrees testing splits to address whether currently available weights can generalize across diverse backgrounds. The dataset provides both within-distribution splits, where each data source is split into train and test, and out-of-distribution splits, where each source is assigned exclusively to either train or test. Using existing public models produced F1 scores ranging from 0.47 to 0.75, highlighting reasonable, but not strong, performance of existing models for many tasks (Tables 2-4).

**Table 2.** Model metrics and results for the TreePoints datasets. The models are ranked by task and F1 score.

|  |  |  |  | Test |  |  |  | Validation |  |  |
|---|---|---|---|---|---|---|---|---|---|---|
| Model | Fine-tuned | Split | Tree Counting (MAE) | Detection Recall | Detection Precision | F1 | Tree Counting (MAE) | Detection Recall | Detection Precision | F1 |
| TreeFormer | Yes | Within-distribution | 19.55 | 0.78 | 0.76 | 0.77 |  | - | - | - |
| TreeFormer | No | Within-distribution | 123.43 | 0.74 | 0.70 | 0.72 | - | - | - | - |
| SAM3 | No | Within-distribution | 40.60 | 0.70 | 0.61 | 0.65 | - | - | - | - |
| CenterNet | Yes | Within-distribution | 41.8 | 0.69 | 0.64 | 0.66 | - | - | - | - |
| TreeFormer | No | Out-of-distribution | 17.2 | 0.84 | 0.78 | 0.81 | 46.9 | 0.35 | 0.94 | 0.51 |
| TreeFormer | Yes | Out-of-distribution | 50.1 | 0.83 | 0.80 | 0.81 | 74.3 | 0.37 | 0.94 | 0.53 |
| SAM3 | No | Out-of-distribution | 37.2 | 0.69 | 0.67 | 0.68 | 59.0 | 0.43 | 0.79 | 0.56 |
| CenterNet | Yes | Out-of-distribution | 83.4 | 0.61 | 0.61 | 0.61 | 248.7 | 0.48 | 0.73 | 0.58 |

Fine-tuning improved within-distribution performance across all three geometries (F1 +0.05 points, +0.15 boxes, +0.16 polygons), and improved out-of-distribution performance for boxes and polygons, but not for points (Tables 2-4). Visual inspection showed that fine-tuned models were able to better differentiate tree boundaries, as well as suppress non-tree false positives in urban areas (Figure 3). As expected, this fine-tuning was more helpful for the within-distribution split where the models could learn about the specifics of the imagery at each site, than for the out-of-distribution split (F1 increase range 0.08-0.09). Performance on a high quality validation set derived from terrestrial LiDAR showed that across geometries performance was lower on the validation data than the test data, with this drop present in both fine-tuned models and in existing public models. This suggests two distinct sources of bias; the specifics of the validation dataset, associated with the difference between the training and validation domain, and hyperparameter overfitting, associated with the additional decrease in the fine-tuned models.

**Figure 3.**
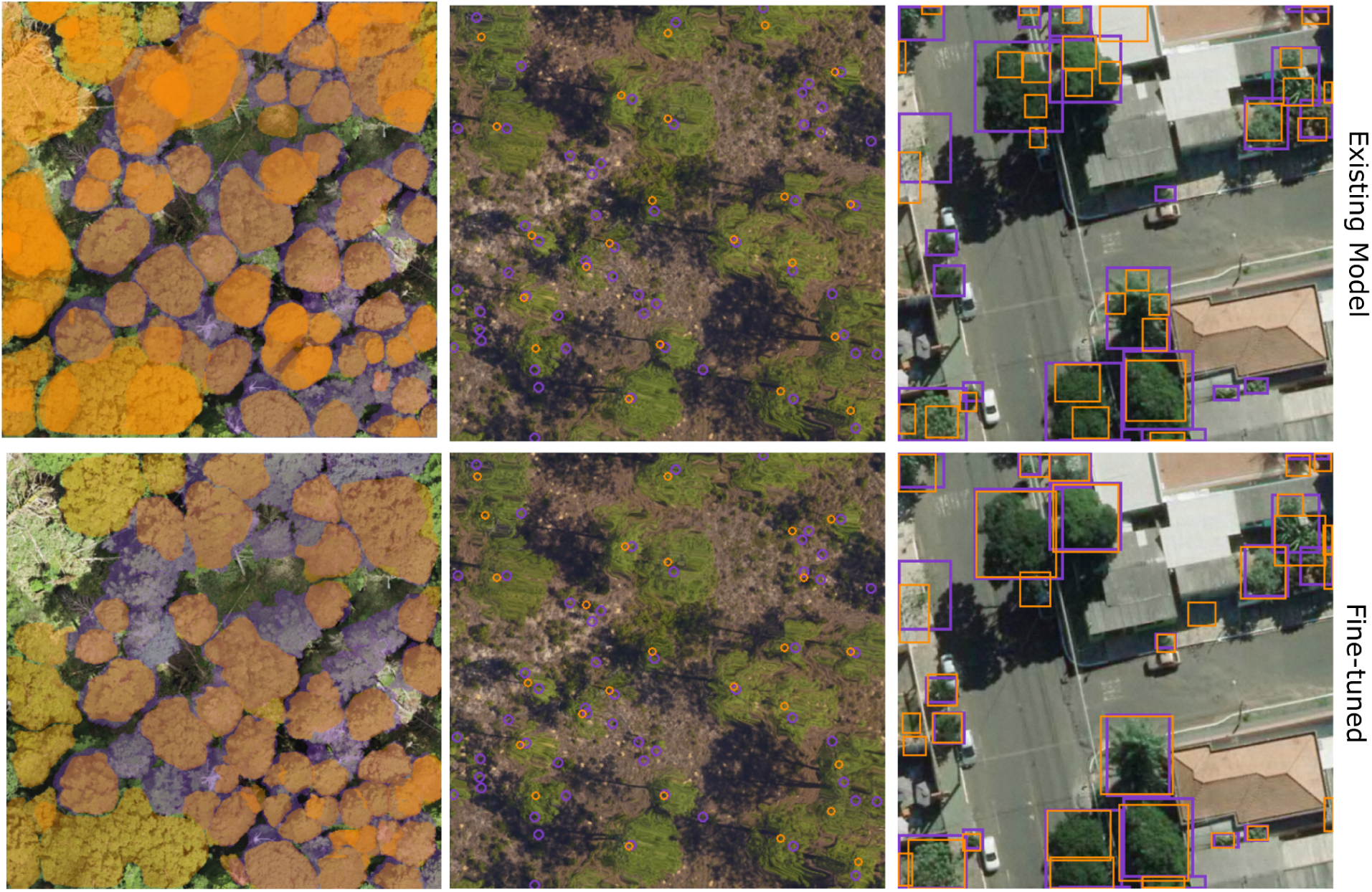
Example within-distribution evaluation image for polygon, point and box datasets from Takeshige et al. 2025, Johnson et al. 2021 and Zamboni et al. 2021. Each dataset is shown with an existing model (DetectTree2, TreeFormer, DeepForest NEON tree detector) and a model fine-tuned on the MillionTrees datasets. For each image the evaluation split with predicted tree detections in orange, and ground truth in purple.

In addition to models specifically designed for detecting and delineating trees we evaluated the performance of the Segment Anything Model 3 (SAM3) foundation model. SAM3 showed variable performance across geometries and datasets. For the point tasks it performed well, but below the bespoke tree detection models for the within and out-of-distribution test tasks (Table 2). For the box tasks it had the highest overall performance for existing public models, but exhibited highly variable performance among data sources, with high scores for some datasets, such as the ones in the out-of-distribution split, and low scores on others including several smaller datasets in the within-distribution split (Table 3, Figure 4). Compared to the bespoke tree detection models, SAM3 had more consistent performance between test and validation datasets, highlighting a lack of hyperparameters and specific tuning to resolution or tree density.

**Figure 4.**
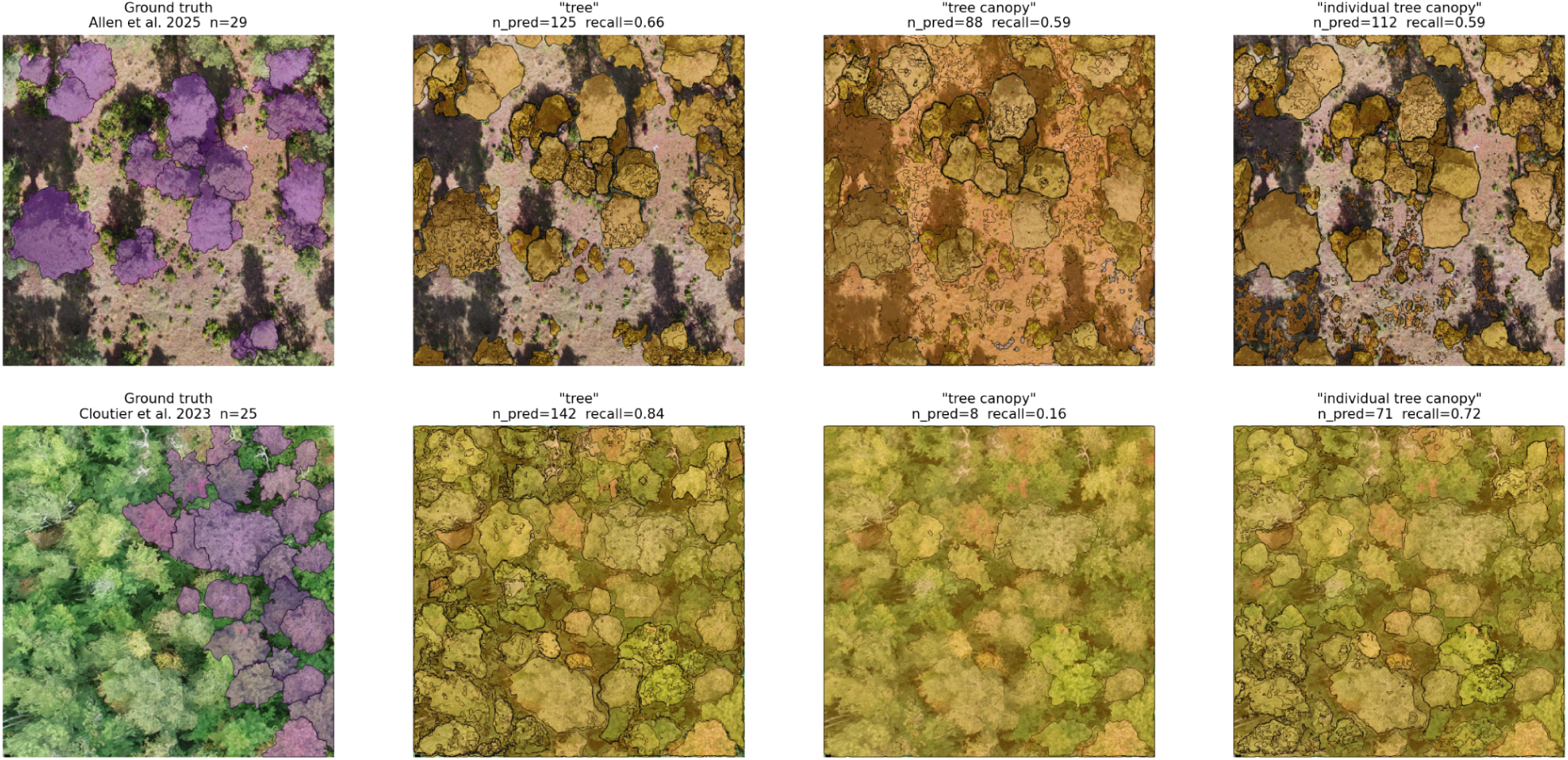
The effect of semantic concept and image background on the quality of SAM3 predictions. The ground truth image (far left) is shown compared to three SAM3 prompts, highlighting the strong variability in performance in foundation models for tree detections. We found that ‘individual tree canopy’ was the most consistent prompt to delineate visible canopy crowns. However, in images with less sharp crown boundaries, the image is flooded with overlapping predictions that lack well-defined edges (second row).

**Table 3.** Model metrics and results for TreeBoxes dataset for the test and validation splits. Models are ranked by task and F1 score.

|  |  |  |  | Test |  |  | Validation |  |  |  |  |
|---|---|---|---|---|---|---|---|---|---|---|---|
| Model | Fine-tuned | Split type | Tree Counting (MAE) | Detection Recall | Detection Precision | F1 | Detection Recall | Detection Precision | F1 | Crown Segmentation AP@40 | Crown Segmentation AP@60 |
| CanopyRS | Yes* | Within-distribution | 12.7 | 0.71 | 0.76 | 0.74 | - | - | - | - | - |
| DeepForest | Yes | Within-distribution | 11.62 | 0.68 | 0.74 | 0.71 | - | - | - | - | - |
| SAM3 | No | Within-distribution | 37.0 | 0.66 | 0.48 | 0.55 | - | - | - | - | - |
| DeepForest | No | Within-distribution | 11.9 | 0.41 | 0.64 | 0.50 | - | - | - | - | - |
| DeepForest | Yes | Out-of-distribution | 13.2 | 0.70 | 0.83 | 0.76 | 0.39 | 0.89 | 0.53 | 0.28 | 0.12 |
| SAM3 | No | Out-of-distribution | 40.2 | 0.72 | 0.58 | 0.64 | 0.57 | 0.60 | 0.58 | 0.41 | 0.20 |
| DeepForest | No | Out-of-distribution | 13.1 | 0.54 | 0.78 | 0.64 | 0.29 | 0.97 | 0.44 | 0.13 | 0.01 |

\*The CanopyRS model was not explicitly fine-tuned on MillionTrees, but it was trained on many of the same sources that are present in the MillionTrees TreeBoxes dataset, as such its scores do not constitute zero-shot performance on this dataset.

For individual geometries, we found a general pattern of F1 scores between 0.5 and 0.7, showing reasonable generalization from training to test, but decline in performance on validation data. The point geometry splits showed strong precision and reasonable recall scores across backgrounds and forest type with the TreeFormer model performing highest on the within-distribution and out-of-distribution splits, but the CenterNet model and SAM3 performing best on the validation data. This complex result comes from several sources; the point detection threshold of within 4m of a ground truth point proved challenging for many of the coarser resolution datasets, which likely reflects both the inherent challenge of predicting a tree center from the airborne perspective, as well as the geospatial error in collecting tree stem locations.

The TreeFormer model in particular was sensitive to hyperparameter and overfitting, with strong fine-tuning within-distribution scores compared to the pretrained model, but these models did not generalize well to the out-of-distribution task. The box geometry results scored in a similar range to point geometry, with the top performing models in the low 70% F1. Best fitting models were CanopyRS (fine-tuned, within-distribution), DeepForest (fine-tuned; out-of-distribution), and SAM3 (existing model).The polygon geometry results show higher precision and lower recall compared to the more even precision-recall tradeoff for the box geometry. Best fitting models were CanopyRS (fine-tuned, within-distribution), SAM3 (existing model, within-distribution), Mask R-CNN (fine-tuned; out-of-distribution), and DetectTree2 (existing model, out-of-distribution). Note that not all models are used for all combinations of fine-tuning and split type. This reflects our focus on determining the general state of tree detection and segmentation, not the best single model, combined with technical and data limitations that do not allow all models to be included in all combinations.

In the cross-geometry task, the model uses only MillionTrees point data to predict crown polygons (Figure 5). We used the fine-tuned TreeFormer model to detect tree centroids and SAM2, a general segmentation model prompted by input geometry, to provide an example of a workflow to address this task. While the F1 score of 0.54 is lower than the best performance on the out-of-distribution polygon task that shares the same test data (Table 4), it is still high enough to be useful given the faster training and annotation times for point data combined with the increasing generality of foundation segmentation models.

**Figure 5.**
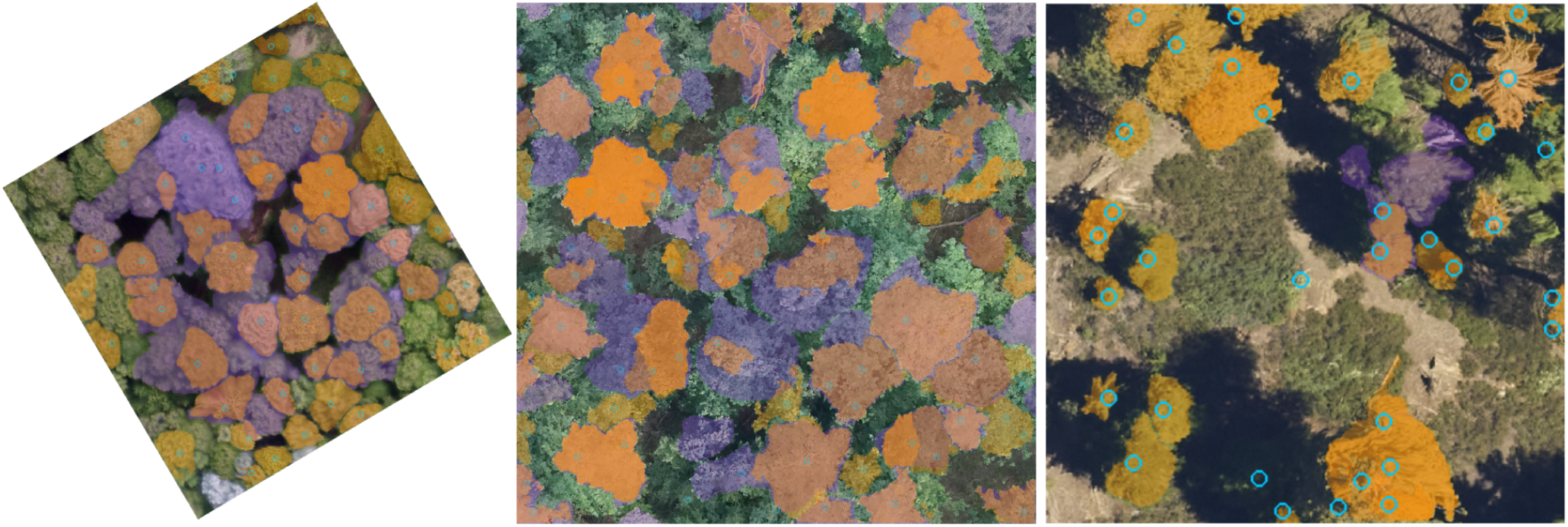
Example images from the cross-geometric prediction task. Point detection using the fine-tuned TreeFormer architecture was first used to predict tree locations, which are then passed to Segment Anything Model 2 (SAM2) to be used as prompts for crown polygon segmentation. Orange polygons are the predicted crowns from the workflow compared to ground truth polygons in purple.

**Table 4.** Model metrics and results for the TreePolygons datasets. Models are ranked by task and test F1.

|  |  |  | Test |  |  | Validation |  |  |  |  |
|---|---|---|---|---|---|---|---|---|---|---|
| Model | Fine-tuned | Split type | Detection Recall | Detection Precision | F1 | Detection Recall | Detection Precision | F1 | Crown Segmentation (AP@40) | Crown Segmentation (AP@60) |
| CanopyRS | Yes* | Within-distribution | 0.73 | 0.87 | 0.79 | - | - | - | - | - |
| Mask R-CNN | Yes | Within-distribution | 0.64 | 0.91 | 0.75 | - | - | - | - | - |
| SAM3 | No | Within-distribution | 0.56 | 0.61 | 0.59 | - | - | - | - | - |
| DetectTree2 | No | Within-distribution | 0.53 | 0.58 | 0.55 | - | - | - | - | - |
| DetectTree2 | No | Out-of-distribution | 0.62 | 0.64 | 0.63 | 0.50 | 0.63 | 0.56 | 0.35 | 0.14 |
| Mask R-CNN | Yes | Out-of-distribution | 0.43 | 0.94 | 0.59 | 0.43 | 0.97 | 0.59 | 0.29 | 0.11 |
| SAM3 | No | Out-of-distribution | 0.56 | 0.67 | 0.61 | 0.51 | 0.58 | 0.55 | 0.34 | 0.14 |
| TreeFormer + SAM2 | No | Cross-geometry | 0.35 | 0.79 | 0.49 | 0.27 | 0.72 | 0.39 | 0.13 | 0.03 |

\*The CanopyRS model was not explicitly fine-tuned on MillionTrees, but it was trained on many sources that are present in the MillionTrees TreePolygons dataset, as such its scores do not constitute zero-shot performance on this dataset.

## Discussion

The MillionTrees benchmark serves as a foundation for airborne tree detection in RGB imagery. This task is highly diverse, with differences among applications in forest type, tree density, tree size, camera resolution, acquisition altitude and illumination conditions. General benchmarks that span these characteristics are crucial for developing models that generalize well to novel contexts and measuring progress towards that goal. Evaluating existing pre-trained weights commonly used for RGB tree detection on this benchmark shows that current models, trained on narrower datasets, remain below state-of-the-art performance in other areas of computer vision and are generally not yet capable of being effectively applied for zero-shot predictions in novel conditions. While we do not know the exact requirement of downstream applications for tree recall and precision, our expectation is that current models will not be sufficient in capturing fine-scaled forest dynamics and applications to ecosystem ecology and silviculture without training on local data and additional investment in field validated ground truth.

The training data in this benchmark can be incorporated into existing algorithms to produce significant improvements in performance. This was not a given, since the datasets contain many different resolutions, annotation protocols and levels of completeness, and it was unknown whether large networks, which are prone to overfitting, had sufficient flexibility to accommodate diverse train and test sources at once. For example, in the within-distribution box tasks, models had F1 scores around 0.5 for existing model weights and between 0.62 and 0.72 for fine-tuned detection, despite coming from different training sources, model architectures and training regimes (Table 3). Our experiments show that fine-tuning on MillionTrees training data improves model performance scores across a range of conditions and highlights three takeaways: 1) current open-source pretrained weights cannot generalize to global scenes that vary in background, resolution, image size, and environment; 2) the MillionTrees dataset can improve these models by large-scale fine-tuning for general airborne tree models; 3) there is still significant room for improvement in the performance of current models even after fine-tuning on this large and diverse data, suggesting a need for ongoing innovations in data collection, model architecture and workflows.

In addition to improvements in the within-distribution split there was an ∼10% absolute improvement for the out-of-distribution split for box and polygon tasks, suggesting that there is significant benefit to training on this broad benchmark for improving the underlying feature representation of trees and therefore producing more generalizable models for unseen sources. This result was not consistent in the Point task, where TreeFormer model was the best performing in the within-distribution test dataset, but did not learn to improve general features for the out-of-distribution dataset within both pretrained and fine-tuned models having similar performance. The lack of continuity in results among geometric tasks underlines a key future need for a single model architecture or training workflow that can harness features from all three geometric sources and be customized to produce the desired geometric result based on the ecological application. Our initial foray into this goal used the large scale weakly labeled box pretraining data to improve the point model for the within-distribution, but not out-of-distribution tasks for points, and were marginally better when co-trained alongside the supervised data for the box and polygon model tasks (Table 5). Combining annotation formats in a flexible framework that allows simultaneous learning on tree centers, crown location and boundaries will be key in capturing the range of tree conditions and acquisition conditions at a global scale.

**Table 5.** Box task performance trained from ImageNet weights with optional inclusion of weak labels. To illustrate the utility of the weakly labeled data sources, we compared the performance of a standard box model with and without inclusion on these unsupervised data for both within-distribution and out-of-distribution tasks for the box geometry. To mimic the behavior of a typical downstream user, the ‘non-pretrained’ model still uses the generic ImageNet-trained weights, rather than random initialized weights.

| TreePoints |  |  |  |  |  |  |
|---|---|---|---|---|---|---|
| Model | Weak Labels | Split | MAE | Recall | Mask-aware Precision | F1 |
| TreeFormer | ✓ | Within-distribution | 28.45 | 0.77 | 0.85 | 0.81 |
| TreeFormer | ✗ | Within-distribution | 66.99 | 0.85 | 0.66 | 0.74 |
| TreeFormer | ✓ | Out-of-distribution | 52.52 | 0.81 | 0.79 | 0.81 |
| TreeFormer | ✗ | Out-of-distribution | 29.42 | 0.86 | 0.79 | 0.82 |
| TreeBoxes |  |  |  |  |  |  |
| Model | Weak Labels | Split | MAE | Recall | Mask-aware Precision | F1 |
| RetinaNet | ✓ | Within-distribution | 12.5 | 0.68 | 0.68 | 0.68 |
| RetinaNet | ✗ | Within-distribution | 11.6 | 0.68 | 0.74 | 0.71 |
| RetinaNet | ✓ | Out-of-distribution | 15.5 | 0.69 | 0.78 | 0.73 |
| RetinaNet | ✗ | Out-of-distribution | 13.2 | 0.70 | 0.83 | 0.76 |
| TreePolygons |  |  |  |  |  |  |
| Mask-RCNN | ✓ | Within-distribution |  | 0.76 | 0.79 | 077 |
| Mask-RCNN | ✗ | Within-distribution |  | 0.65 | 0.91 | 0.76 |
| Mask-RCNN | ✓ | Out-of-distribution |  | 0.69 | 0.79 | 0.73 |
| Mask-RCNN | ✗ | Out-of-distribution |  | 0.43 | 0.94 | 0.59 |

Traditional computer vision benchmarks are designed to have ideal data, with completely annotated imagery and minimal annotation uncertainty, with the aim of focusing on improving model architecture or training regimes (Koh et al., 2021). One of the central challenges to general solutions for airborne tree detection is that this ideal data has 1) large numbers of annotations; 2) sufficiently diverse sets of annotations to create robust models; 3) images where all visible trees are annotated; and 4) annotations that represent individual trees as accurately as possible, either through field-based confirmation or multi-sensor agreement to reduce annotation uncertainty. The final requirement is particularly relevant for trees since they are often only partially visible in overhead RGB imagery, with overlapping crowns among neighbors, making it difficult to annotate the precise location of one tree versus the surrounding trees (Baudchon et al., 2026, Allen et al., 2025). Accomplishing all four of these components is typically not possible within a single dataset due to tradeoffs in the time and cost required to annotate a single image versus the number and diversity of images that can be annotated (Mishra 2026). To address this challenge we take an inclusive approach, incorporating the full range of annotation approaches to provide the most data possible, providing users with easy ways to subset to the particular annotation types they want to use, and prioritizing the inclusion of more accurate, costly and therefore limited data in the evaluation data splits. This does mean that a significant portion of datasets are incompletely annotated. We included an annotation mask with each image to allow the use of incompletely annotated images while reducing the risk of models suppressing true positive truths due to unannotated portions of training data. Existing models, for example Ball et al. 2023, used tiles with up to 60% unannotated crowns, and more analysis is needed to determine the optimal way to use the annotation masks to maximize the utility of incomplete datasets without negatively impacting model performance. For example, directly using the annotation completeness mask can allow models to balance feature learning and suppression of false-backgrounds using additional loss functions (e.g. Zhang et al., 2020).

The limitations in annotation directly inform which ecological tasks are most appropriate for airborne tree detection. Tasks that are highly sensitive to small trees or measurements of crown area are the most difficult with current data and model limitations. Applications of tree counting are further complicated by subcanopy trees not being visible in airborne imagery, which can result in mismatches between the field-countable and airborne observable stems (e.g. Davies et al. 2021). There is no well-defined protocol for deciding which trees to include as ground truth and how to associate forest plot metadata with top-down sensor pixel data. Due to tree lean and competition for light resources, the point centers taken at stem coordinates common in forest surveys are often not directly under the crowns visible from the airborne perspective (Graves et al. 2018). Young et al. (2022) demonstrated strong performance of a greedy algorithm for ground-to-aerial tree matching based on proximity of similarly sized trees when allowing greater distance tolerance for taller trees. In situations where tree height and airborne LiDAR data are present, establishing subdominant crowns by comparing heights and surrounding canopy height (e.g. Young et al. 2022) is useful, but not a foolproof way of distinguishing subcanopy data (Weinstein et al. 2024). This challenge can also lead to an underestimation of subcanopy resources that may impact downstream applications, especially for measurements of standing wood, silviculture or ecosystem services. This limitation has been noted for other airborne sensors as well (Cao et al. 2023). While there have been statistical attempts to infer subcanopy structure from airborne perspectives (Duncanson et al. 2014, Fischer et al. 2020), the focus of the MillionTrees benchmark is to improve canopy tree observation. Given our results, there are still substantial areas for improvement for a flexible global model of canopy tree detection before the inclusion of additional complexity in estimating unseen forest structure.

The process of compiling this benchmark highlighted that the highest quality evaluation data is laborious to collect and accounts for a small proportion of available datasets. Top-down views of dense canopies make it difficult to distinguish among neighboring trees. Bottom-up views can aid by comparing the number of stems visible from under the canopy, but the boundaries of trees are difficult to estimate in multi-layered canopies. Simultaneous views from above and below, such as co-registered terrestrial LiDAR and RGB orthomosaics increase the confidence for validation accuracy. Terrestrial LiDAR algorithms for tree segmentation can help speed up the annotation process (Devereux et al. 2026, Xiang et al. 2025, Henrich et al., 2024, Wielgosz et al. 2026), but this remains a significant hurdle as manual corrections are generally needed (Cherlet et al. 2026). For a portion of the MillionTrees validation data in Liu et al. 2024, human review was needed to manually verify and join the oversplitting of crowns in the initial TLS segmentation. Human delineation of trees directly onto remote sensing imagery while in the field observing the trees from underneath can similarly leverage views from both above and below the trees to yield more accurate labels (Graves et al. 2018). Extending these kinds of datasets to include complex tropical forests, standardizing the exclusion of subcanopy trees, and connecting these data to traditional forestry plots will help create a gold standard for airborne RGB tree segmentation. However it is unlikely that there will ever be enough of this data to train models due to the time and expense of collecting this type of data. We envision a future where models are trained first on weak labels, then on large image-annotated or field-aligned datasets, and are finally fine-tuned or evaluated using as many of the highest quality annotations as possible.

To leverage our most precisely annotated datasets we reserved these datasets to serve as a validation dataset. Validation datasets are run after test metrics are collected and are based on terrestrial LiDAR-derived annotations (Figure 2). This dataset should not be evaluated at any point during model training or fine tuning and only used for evaluation once all training has been completely finalized. This is important for: 1) assessing whether the hyperparameters used for evaluating test accuracy can generalize to new datasets; and 2) ensuring the model performance metrics reflect the ability to detect individual trees as opposed to the proportion of trees that a human can see within the image in suboptimal circumstances (e.g., when annotating only using airborne imagery; Allen et al. 2025). The difference between human-based image annotation and the true ground truth count can be large, and the difference was tolerable during the initial phases of tree detection research, but as we move towards applications with important real-world consequences, the limitations in evaluation rigor undermine confidence in these models and will hinder widespread adoption and credibility. In recognition of the fine-grained nature of terrestrial LiDAR data in delineating crown boundaries, we used the Average Precision (AP) metric at two intersection-over-union levels to describe the ability for models to perform detailed crown segmentation. We find that models generally describe locations of crowns at the coarser AP@40, but rarely can reproduce crown boundaries at AP@60. Especially for small crowns and coarse resolution data, a prediction can only be offset from ground truth by a small number of pixels to be considered true positive (Figure S1). The AP@60 for several models is below 0.2, suggesting that current models are not suitable for ecological analysis that need highly precise measures of crown area.

The reduced validation scores compared to test scores suggests that hyperparameter selection, especially across resolutions and tree sizes, is especially important for model generalization. In particular, the TreeFormer point model significantly dropped on the validation dataset, suggesting poor ability to generalize between human-verified ground truth and TLS-derived ground truth with a high sensitivity to hyperparameter selection (Figure 6). The same was true to a lesser extent for the DeepForest box model, which declined from test to validation scores, but inspection of these results suggested it was the minimum cutoff score and poor confidence calibration, eliminating many reasonable predictions in the validation dataset compared to the test sources. Large neural networks are notoriously poorly calibrated, and the confidence scores of the model should not be used directly without further review in downstream applications (Guo et al. 2017, Ye et al. 2023).

**Figure 6.**
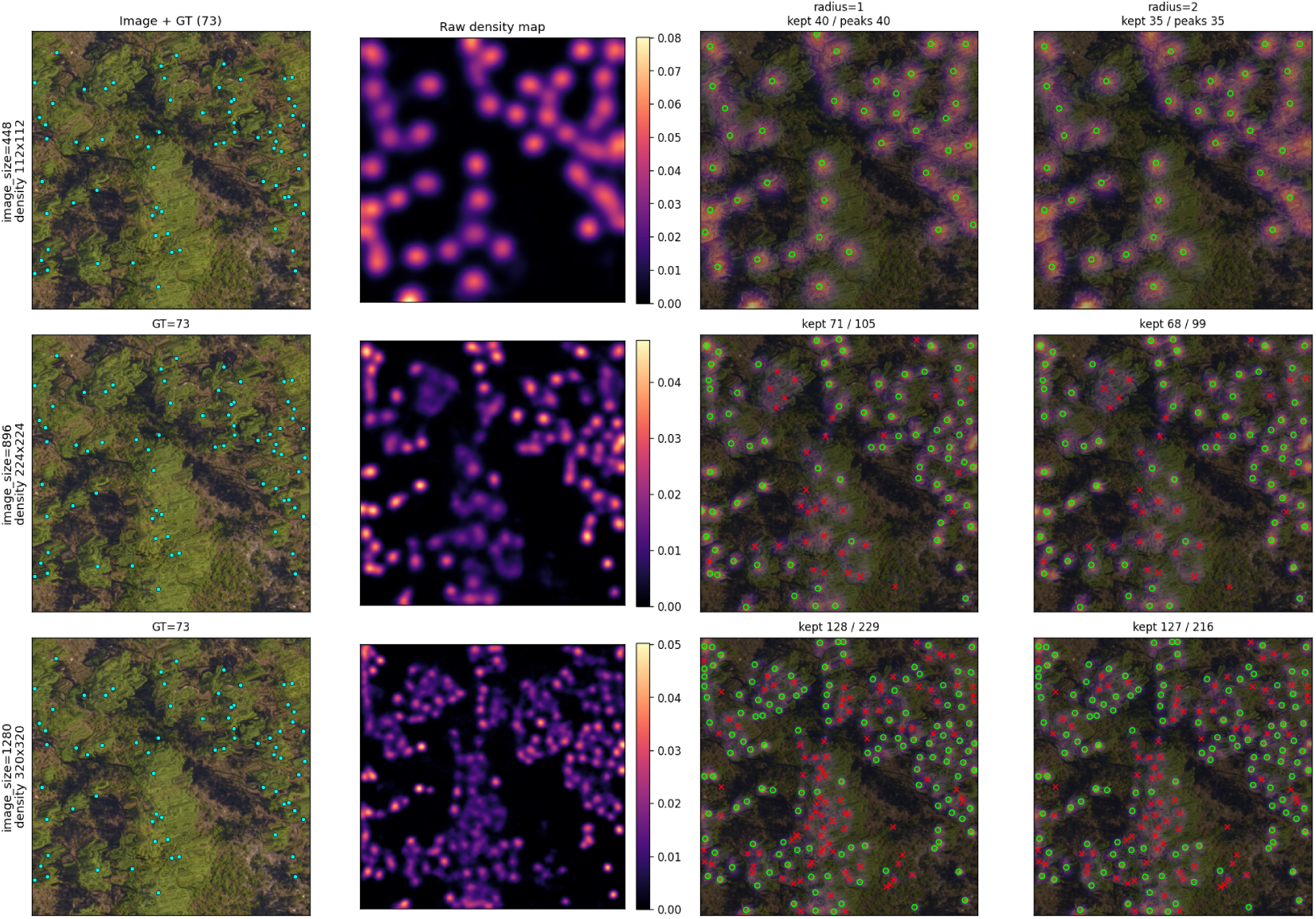
Impact of hyperparameters across spatial resolutions for the TreeFormer point model. The same model weights are used in each panel, but the image size and radius hyperparameter are varied. This illustrates that while the underlying features may produce convincing tree counts and locations, the choice of post-training workflows has significant effects on model accuracy. The change in prediction counts with image size highlights that models may not be robust to transformations in scale and produce widely different prediction counts based on image rescaling and resolution.

Combining all available annotation approaches allows the MillionTrees dataset to stimulate research into model robustness across not only ecosystems, but also across image acquisition variables and hyperparameters including image size, focal view and pixel resolution. During the process of evaluating existing models we found that current models are often highly sensitive to these acquisition variables, hyperparameters, and as a result could perform differentially even on data sources with very similar appearances or on the same data source but varying aspects of the image presentation (e.g., Figure 4). For example, the performance of the Segment Anything Model 3 (SAM3) foundation model was brittle, with excellent performance on some sources and then relatively poor performance on sources that appear visually similar (Figure 4). Overall, SAM3 performed better on point and box tasks and higher resolution data, but less well on polygon tasks and coarser resolution data. Likewise, changing the hyperparameters that govern the size and representation of image features had a large effect on the ability of models to generalize across resolutions, especially for point datasets. Even when trained on sources that vary in these aspects, we found that the models still produced substantially different predictions during inference depending on the selected hyperparameters. Size-based data augmentation during training is standard practice for many of the models tested, but appears to be insufficient to address these changes in object size and variation. The sensitivity to hyperparameters is underappreciated and more work is needed both to train models that are invariant to differences in image size and resolution, and to understand whether fine-tuning is primarily learning new image features or rescaling existing features for the target datasets.

The MillionTrees benchmark helps coalesce a wide body of work into a single set of sources and evaluation scores that can be tracked over time. The flexible nature of the package will allow new data sources and tasks to be added while maintaining the reproducibility of previous scores. There are a diverse set of potential future tasks including generalizing across drone and plane-based sensors to satellite resolutions, model calibration scores to assess the utility of using downstream confidence scores in ecological applications, as well as specific fine-tuning tasks that assess the number of annotations are needed to achieve full-dataset performance. Additional validation datasets will facilitate robust examination on the applications and bias in airborne tree detection. A combination of novel architectures, data selection techniques, and pretraining strategies will improve both within-distribution and out-of-distribution scores to move the open-source community towards a global model of airborne tree detection.

## Materials and Methods

The MillionTrees benchmark focuses on high-resolution (<50 cm) RGB images due to their wide availability and uniform calibration across manufacturers. Datasets were gathered from public repositories, by contacting authors of recent tree detection research articles, and by generating new datasets through both annotation and combining existing ground-based data with public orthoimagery. Datasets represent the current state of annotation for tree detection, and as such cover a range of backgrounds, include a variety of annotation styles and completeness, and incorporate annotations generated from airborne and terrestrial LiDAR.

### Annotation Approaches

There are three main approaches for creating annotations of individual trees for machine learning models: 1) image-annotation, 2) field-based crown delineation, and 3) combining airborne imagery with existing field stem maps. There are tradeoffs for each method, and many studies use a combination of approaches to balance the time investment of ground surveys, the logistics of collecting sufficient data for model development, and the confidence of field-verified evaluation data in dense canopy environments. The most common strategy is image-annotation, in which a human draws the location of each tree directly on the airborne image, optionally alongside additional co-registered remote sensing data (e.g., HSI, LiDAR in Weinstein et al. 2020, infrared in Schutte et al. 2025). The benefits of this approach are the scale of annotation and the clear connection between the annotation and the airborne viewpoint. However, because the primary view of the trees is from above these annotations can combine multiple nearby trees into a single annotation or split a single large tree into multiple annotations. Field-based crown delineation covers a range of approaches including drawing the crowns on existing imagery while standing under the canopy (Graves et al., 2018), field verified marker-assisted views from photogrammetry (Cloutier et al. 2023), and high-precision ground-based spatial survey data co-registered to imagery (Open Forest Observatory 2026). These methods are less likely to lump or split individual trees, but it is much more time consuming, leading to smaller datasets and incompletely annotated scenes. Crown delineation can also be aided by existing stem maps from forestry surveys (e.g. Johnson et al. 2021), municipal urban forestry departments (e.g. Beery et al. 2022) and urban tree programs (Velasquez-Camacho et al. 2023), that can be connected to airborne data to capture tree locations at large extents and used to inform delineation in complex forested scenes. However, these surveys tend to be patchy, leading to many visible trees in airborne imagery that were not included in survey areas or protocols. In addition, hand-held GPS units often cannot reflect the required geospatial accuracy in the subcanopy, and ground-based stations can have difficulty acquiring sufficiently precise readings in forested environments (Kattenborn et al. 2021).

In closed canopy or densely planted plantations scenes, annotation is time consuming and it is common practice to incompletely annotate each image in order to prioritize greater confidence in individual annotations and to generate a sufficiently large and diverse set of images. This creates challenges for most object detection algorithms, since unannotated portions of the image are considered to not include trees during model training. The degree to which incomplete annotation impacts model performance varies among algorithms (Liu et al. 2023, Ball et al. 2023) and is an important area of application for this benchmark. To facilitate the careful use or exclusion of incompletely annotated data we used the semantic segmentation model from Veitch-Michaelis et al. (2024) to label each pixel in each image as ‘Tree Canopy’ or ‘Background’ (Figure 9). This data layer is designed to allow users to filter training data based on completeness at a granular level, as well as develop new methods that can be applied to object detection with incomplete data. The segmentation model is designed to aid in removing misalignments between ground truth and airborne imagery, but cannot distinguish among annotation errors in splitting or lumping trees.

There are multiple approaches to quantifying the location and geometry of individual trees in the canopy. We include all three major geometries used to annotate individual trees: 1) points, which mark the center of the crown or the basal stem location with a single x, y coordinate, 2) boxes, which mark the maximum x, y boundaries of the crown with a rectangle; and 3) polygons, which smoothly mark the boundary of trees using polygons with >4 vertices. Raster-based representations of trees were excluded since they are rarely used to natively represent individual tree data.

To overcome the limitations of the commonly available field-based surveys and image-annotation methods, it is possible to collect high-resolution LiDAR datasets of a forest from the ground and airborne imagery in tandem. The detailed 3D LiDAR scan can be used to create high-quality 3D annotations of tree instances. These can be projected onto 2D labels from an aerial perspective, using the highest-point instance label per pixel as the top visible tree. There are several algorithms to automatically perform single-tree segmentation from high-quality LiDAR data using the full 3D information preserved in the scanning data (e.g. Nguyen et al. 2026, Wielgosz et al. 2026). This also has the potential to make lower trees in occluded areas of the aerial imagery visible, that would otherwise be indistinguishable by a human annotator. Also, other common artifacts like tree forks leading to two crowns from one individual, or very homogeneous crown spaces from multiple trees, can be correctly assigned since the trunk structure is visible in such datasets.

### Dataset Structure

To aid rapid development, the MillionTrees benchmark contains easy to use PyTorch dataloaders that yield images and labels in a consistent AI-ready format without the user needing to align and organize data. To facilitate ease of data sharing and mitigate privacy concerns, and therefore maximize the number of data sources in this benchmark, all data have been stripped of geospatial information and additional metadata including crown area, species identity and tree attributes. The source citation associated with every image is provided so that users of the benchmark can directly connect data with upstream location and citations. The dataloaders are divided into three datasets based on annotations in point, box or polygon format. To standardize image sizes before packaging, we tiled images (while retaining their original resolutions) to between 1000-2000 pixels to reduce the memory load of large images, especially from high resolution drone datasets (e.g. Cloutier et al. 2023). In order to create image batches for training, evaluation and inference, large images have been tiled to a smaller size, and are then resized during batch generation. For each dataloader, there are multiple evaluation tasks. Each task separates the datasets into training images and test images such that annotations in train never occur in test. The conditions that determine the source split varies to reflect different applied scenarios (Figure 7).

**Figure 7.**
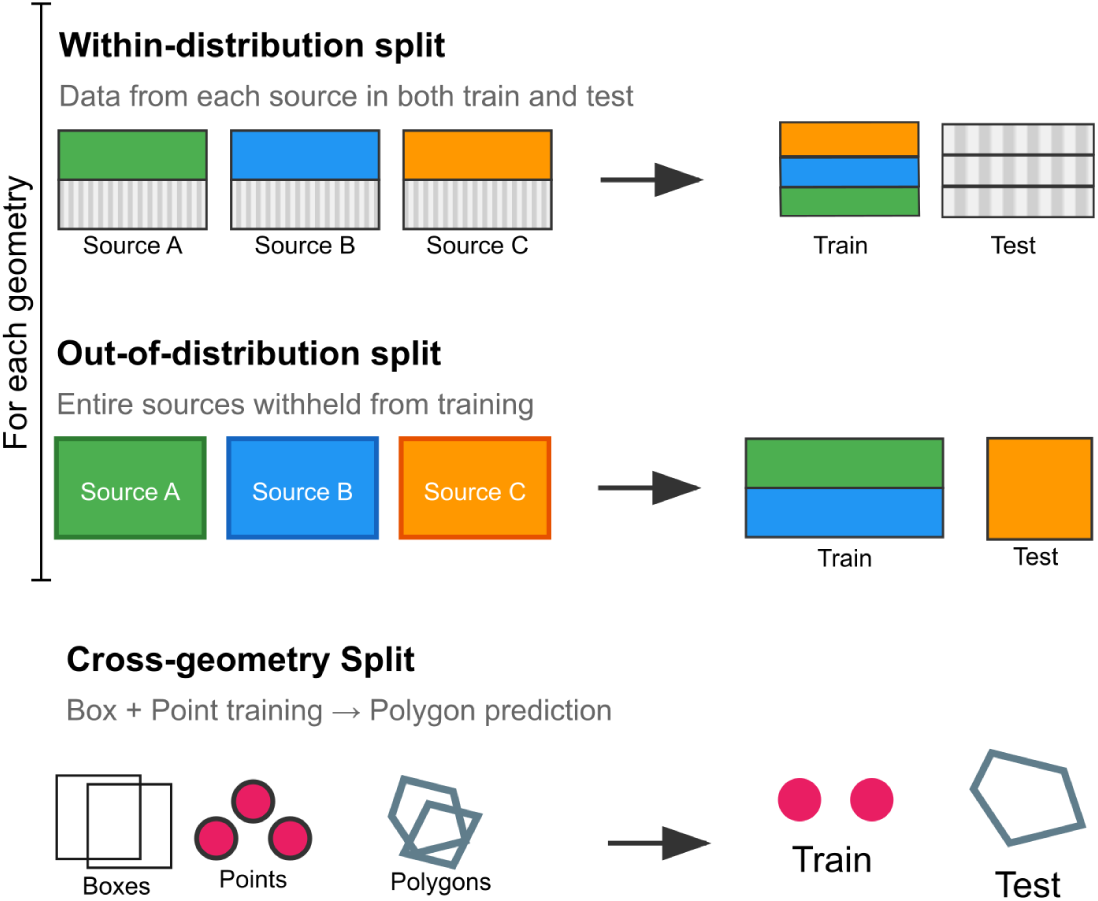
Training and test splits for the MillionTrees tasks. In the within-distribution split, each source is split into train and test to assess the model’s ability to customize to new datasets where annotations are available. In the out-of-distribution split, sources are withheld from training to assess the ability of a model to detect trees in unseen conditions. In the cross-geometry split, points are used to predict polygon tree crowns.

#### Within-distribution split

In the within-distribution task, a portion of the images from each source are split into 90% train and 10% test, with each source occurring in both train and test. Wherever possible we maintained a spatial split, such that image tiles were chosen from different regions of the source imagery. The within-distribution split represents a common use case for applied computer vision in biology. Often the target project has some labels that can be used to fine-tune an existing large pretrained backbone. Our goal with the within-distribution task is to assess model performance given additional training and hyperparameter selection. To get a single metric among test sources, we compute the metrics per-source and then average among sources to highlight the flexibility of proposed models in a range of backgrounds and annotation sizes.

#### Out-of-distribution split

In the out-of-distribution split task, each source is placed into either train or test. This supports evaluating the transferability of existing models to new sources of data, which is a challenging scenario due to the variable nature of tree detection across geographies, background and taxa. We prioritized test sources with field-validation and because of this there are fewer test sources than in the within-distribution task. The number and diversity of sources vary among the point, box and polygon geometry tasks and should not be compared since they may vary in how similar they are to the ecologies present in train sources.

#### Cross-geometry split

The vast majority of benchmarks in ecological computer vision contain a single type of annotation geometry and format. To go beyond these representations of ecological objects, we propose a task to foster creativity and novel models that can flexibly utilize multiple representations during pretraining, training and inference. This flexibility will allow greater inclusion of data sources and generalization to new scenes. This ‘cross-geometry’ task uses tree points for training and polygon predictions as the output. Two polygon sources were chosen for the evaluation data and all point sources are placed in the training set. We chose this combination to mimic the tradeoff in annotation completeness and effort. Polygon inference is often desirable for downstream tasks such as species classification and crown measurements, but is more costly to collect, and therefore less commonly available due to the greater time and effort required to create polygon annotations.

#### Source Datasets

The source datasets are a collection of annotations that vary in geography, background, tree density and annotation protocols. To catalog the datasets, we consider them broadly as ‘image-annotated’, ‘field-collected’ and ‘pseudo-labeled’. These designations represent the tradeoff between annotation completeness, confidence and scale and reflect the ongoing research on the best way to collect sufficient data to train general models for classes at global scale (Table 6). Image-annotation is often used in our sources in forested scenes to maximize the number of tree annotations in dense images (Firoze et al., 2023, Lucas et al., 2024), with scenes from Germany (Troles et al., 2024) and Sweden (Swedish Forest Agency, 2021) designed for tree detection tasks, while others such as Jansen et al. (2023) or Puliti and Astrup (2025) were annotated for tree species classification or tree-top damage detection, and used here for tree detection. Image-annotated datasets also include urban scenes (Amirkolaee et al., 2023; Kwon et al., 2023; Velasquez-Camacho et al., 2023; Zamboni et al., 2021), where field validation of all trees in the image may not be logistically possible due to accessing private property. While annotations were made directly on the images from the airborne perspective, many datasets were accompanied with additional data sources to enhance their accuracy. For example, Velasquez-Camacho et al. 2023 used urban forestry records to correct the locations of airborne annotations and Weinstein et al. 2020 were annotated by simultaneously viewing coregistered RGB, LIDAR, and Hyperspectral imagery.

**Table 6.**
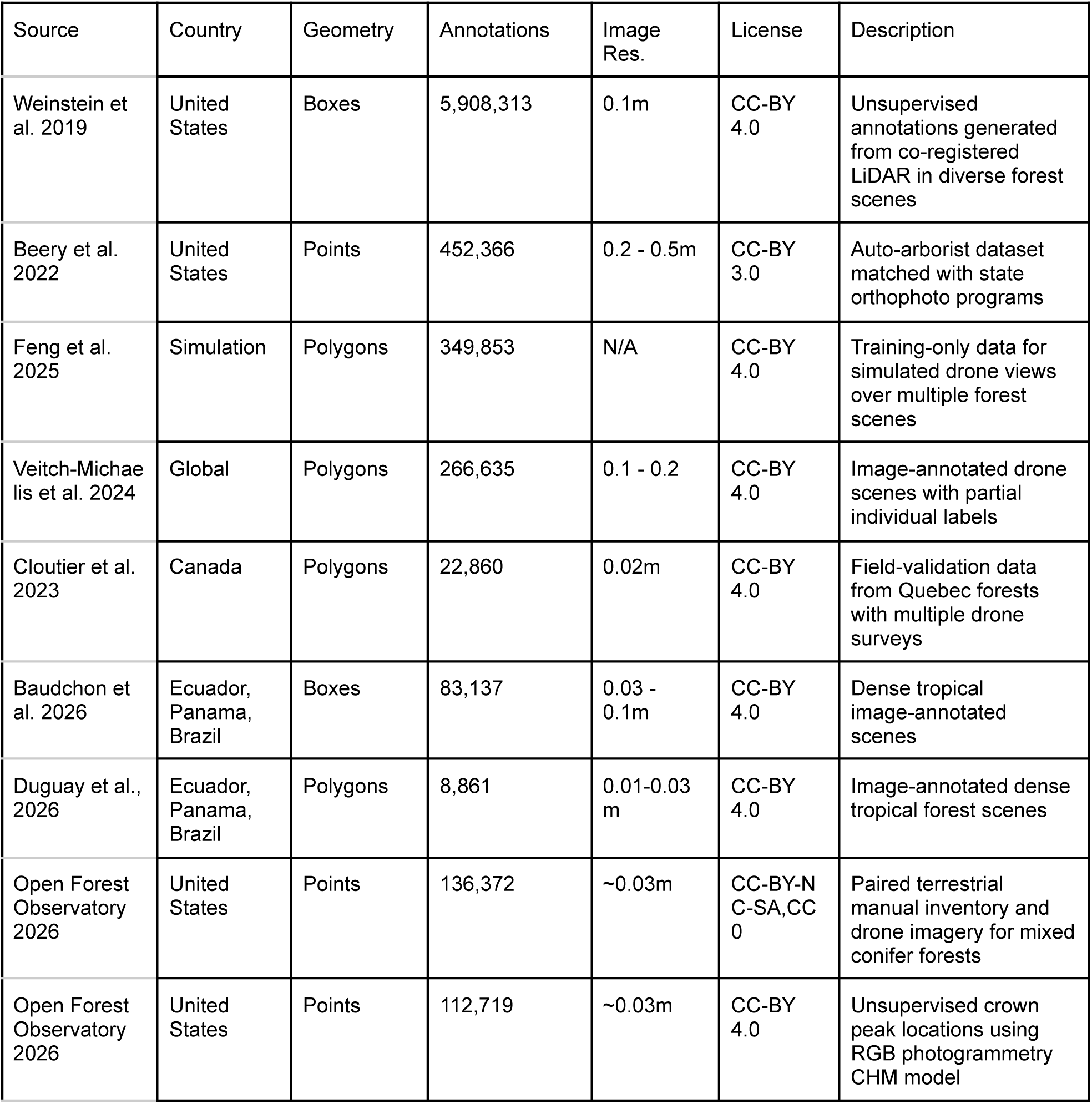
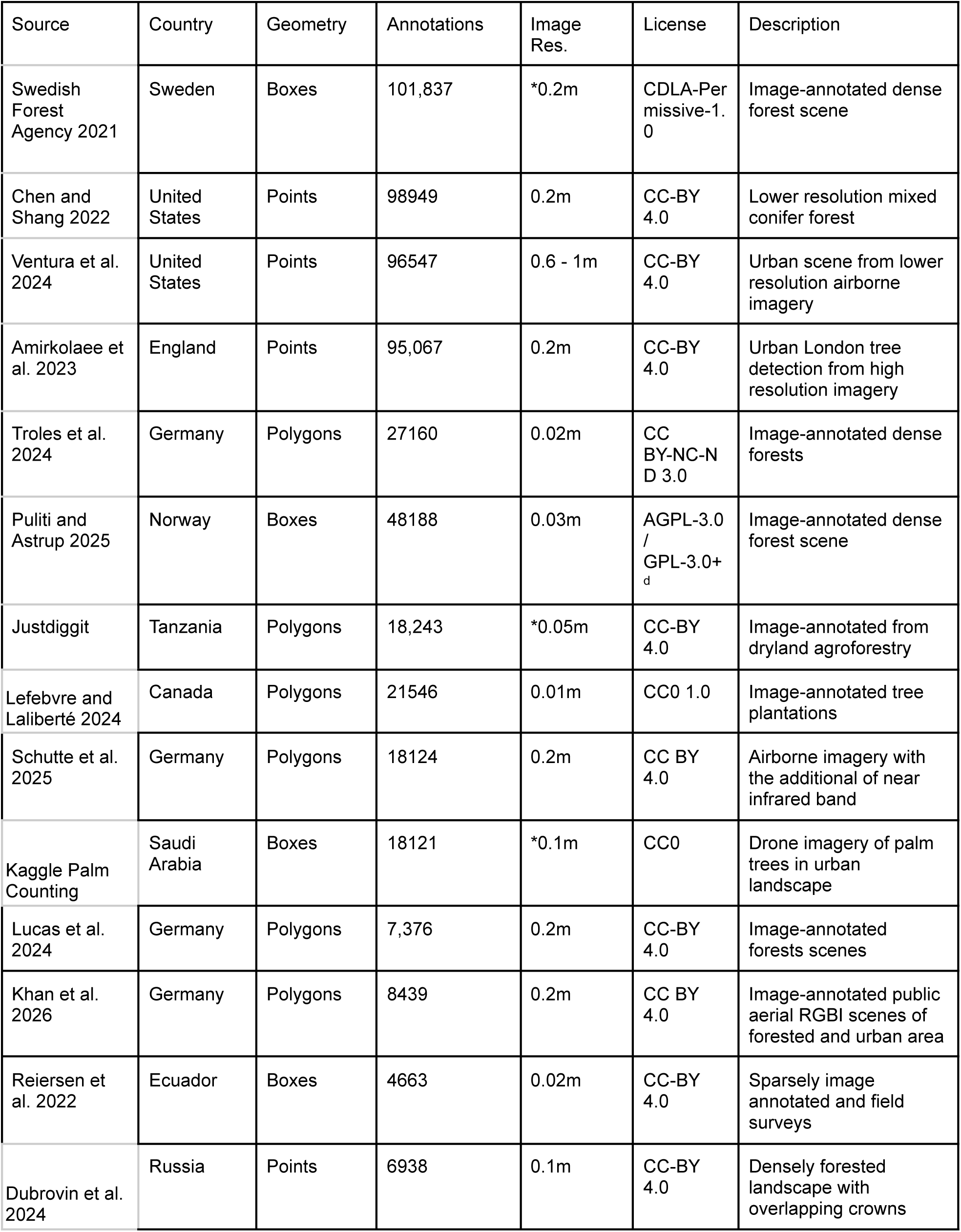
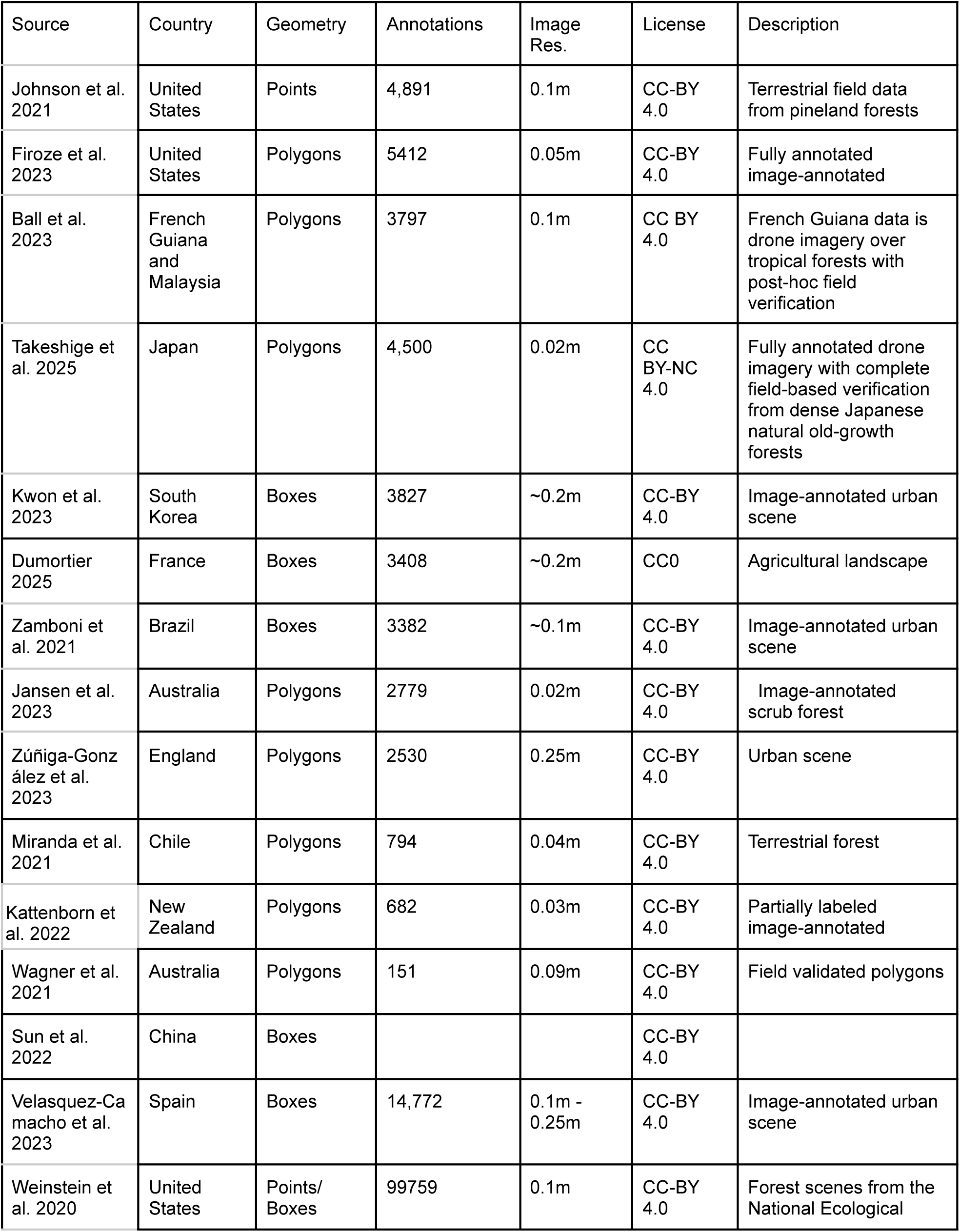
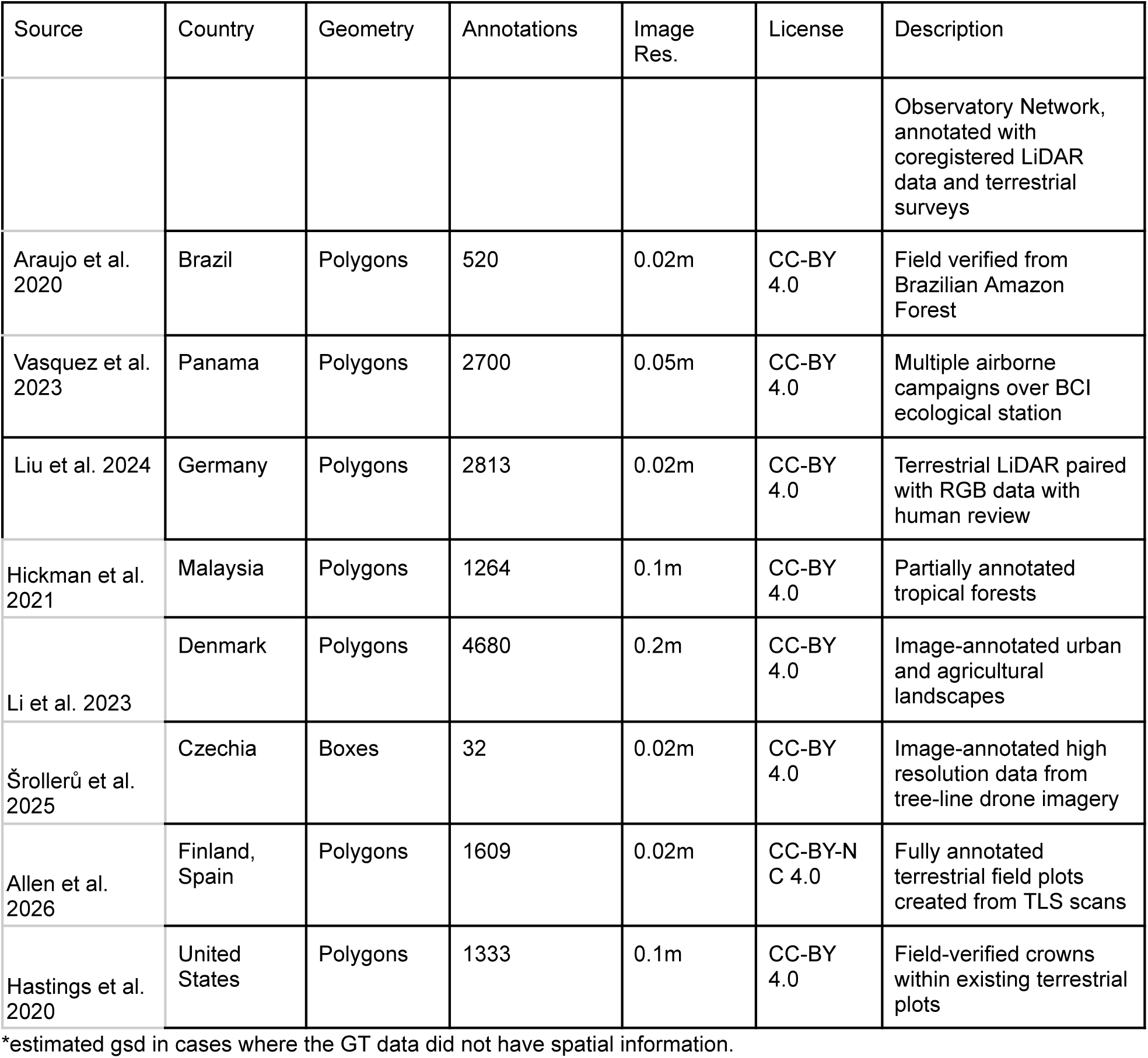
Description of source datasets.

| Source | Country | Geometry | Annotations | Image Res. | License | Description |
|---|---|---|---|---|---|---|
| Weinstein et al. 2019 | United States | Boxes | 5,908,313 | 0.1m | CC-BY 4.0 | Unsupervised annotations generated from co-registered LiDAR in diverse forest scenes |
| Beery et al. 2022 | United States | Points | 452,366 | 0.2 - 0.5m | CC-BY 3.0 | Auto-arborist dataset matched with state orthophoto programs |
| Feng et al. 2025 | Simulation | Polygons | 349,853 | N/A | CC-BY 4.0 | Training-only data for simulated drone views over multiple forest scenes |
| Veitch-Michaelis et al. 2024 | Global | Polygons | 266,635 | 0.1 - 0.2 | CC-BY 4.0 | Image-annotated drone scenes with partial individual labels |
| Cloutier et al. 2023 | Canada | Polygons | 22,860 | 0.02m | CC-BY 4.0 | Field-validation data from Quebec forests with multiple drone surveys |
| Baudchon et al. 2026 | Ecuador, Panama, Brazil | Boxes | 83,137 | 0.03 - 0.1m | CC-BY 4.0 | Dense tropical image-annotated scenes |
| Duguay et al., 2026 | Ecuador, Panama, Brazil | Polygons | 8,861 | 0.01-0.03m | CC-BY 4.0 | Image-annotated dense tropical forest scenes |
| Open Forest Observatory 2026 | United States | Points | 136,372 | ~0.03m | CC-BY-NC-SA,CC0 | Paired terrestrial manual inventory and drone imagery for mixed conifer forests |
| Open Forest Observatory 2026 | United States | Points | 112,719 | ~0.03m | CC-BY 4.0 | Unsupervised crown peak locations using RGB photogrammetry CHM model |
| Swedish Forest Agency 2021 | Sweden | Boxes | 101,837 | *0.2m | CDLA-Permissive-1.0 | Image-annotated dense forest scene |
| Chen and Shang 2022 | United States | Points | 98949 | 0.2m | CC-BY 4.0 | Lower resolution mixed conifer forest |
| Ventura et al. 2024 | United States | Points | 96547 | 0.6 - 1m | CC-BY 4.0 | Urban scene from lower resolution airborne imagery |
| Amirkolaee et al. 2023 | England | Points | 95,067 | 0.2m | CC-BY 4.0 | Urban London tree detection from high resolution imagery |
| Troles et al. 2024 | Germany | Polygons | 27160 | 0.02m | CC BY-NC-ND 3.0 | Image-annotated dense forests |
| Puliti and Astrup 2025 | Norway | Boxes | 48188 | 0.03m | AGPL-3.0 / GPL-3.0+<sub>d</sub> | Image-annotated dense forest scene |
| Justdiggit | Tanzania | Polygons | 18,243 | *0.05m | CC-BY 4.0 | Image-annotated from dryland agroforestry |
| Lefebvre and Laliberté 2024 | Canada | Polygons | 21546 | 0.01m | CC0 1.0 | Image-annotated tree plantations |
| Schutte et al. 2025 | Germany | Polygons | 18124 | 0.2m | CC BY 4.0 | Airborne imagery with the additional of near infrared band |
| Kaggle Palm Counting | Saudi Arabia | Boxes | 18121 | *0.1m | CC0 | Drone imagery of palm trees in urban landscape |
| Lucas et al. 2024 | Germany | Polygons | 7,376 | 0.2m | CC-BY 4.0 | Image-annotated forests scenes |
| Khan et al. 2026 | Germany | Polygons | 8439 | 0.2m | CC BY 4.0 | Image-annotated public aerial RGBI scenes of forested and urban area |
| Reiersen et al. 2022 | Ecuador | Boxes | 4663 | 0.02m | CC-BY 4.0 | Sparsely image annotated and field surveys |
| Dubrovin et al. 2024 | Russia | Points | 6938 | 0.1m | CC-BY 4.0 | Densely forested landscape with overlapping crowns |
| Johnson et al. 2021 | United States | Points | 4,891 | 0.1m | CC-BY 4.0 | Terrestrial field data from pineland forests |
| Firoze et al. 2023 | United States | Polygons | 5412 | 0.05m | CC-BY 4.0 | Fully annotated image-annotated |
| Ball et al. 2023 | French Guiana and Malaysia | Polygons | 3797 | 0.1m | CC BY 4.0 | French Guiana data is drone imagery over tropical forests with post-hoc field verification |
| Takeshige et al. 2025 | Japan | Polygons | 4,500 | 0.02m | CC BY-NC 4.0 | Fully annotated drone imagery with complete field-based verification from dense Japanese natural old-growth forests |
| Kwon et al. 2023 | South Korea | Boxes | 3827 | ~0.2m | CC-BY 4.0 | Image-annotated urban scene |
| Dumortier 2025 | France | Boxes | 3408 | ~0.2m | CC0 | Agricultural landscape |
| Zamboni et al. 2021 | Brazil | Boxes | 3382 | ~0.1m | CC-BY 4.0 | Image-annotated urban scene |
| Jansen et al. 2023 | Australia | Polygons | 2779 | 0.02m | CC-BY 4.0 | Image-annotated scrub forest |
| Zúñiga-González et al. 2023 | England | Polygons | 2530 | 0.25m | CC-BY 4.0 | Urban scene |
| Miranda et al. 2021 | Chile | Polygons | 794 | 0.04m | CC-BY 4.0 | Terrestrial forest |
| Kattenborn et al. 2022 | New Zealand | Polygons | 682 | 0.03m | CC-BY 4.0 | Partially labeled image-annotated |
| Wagner et al. 2021 | Australia | Polygons | 151 | 0.09m | CC-BY 4.0 | Field validated polygons |
| Sun et al. 2022 | China | Boxes |  |  | CC-BY 4.0 |  |
| Velasquez-Camacho et al. 2023 | Spain | Boxes | 14,772 | 0.1m - 0.25m | CC-BY 4.0 | Image-annotated urban scene |
| Weinstein et al. 2020 | United States | Points/ Boxes | 99759 | 0.1m | CC-BY 4.0 | Forest scenes from the National Ecological Observatory Network, annotated with coregistered LiDAR data and terrestrial surveys |
| Araujo et al. 2020 | Brazil | Polygons | 520 | 0.02m | CC-BY 4.0 | Field verified from Brazilian Amazon Forest |
| Vasquez et al. 2023 | Panama | Polygons | 2700 | 0.05m | CC-BY 4.0 | Multiple airborne campaigns over BCI ecological station |
| Liu et al. 2024 | Germany | Polygons | 2813 | 0.02m | CC-BY 4.0 | Terrestrial LiDAR paired with RGB data with human review |
| Hickman et al. 2021 | Malaysia | Polygons | 1264 | 0.1m | CC-BY 4.0 | Partially annotated tropical forests |
| Li et al. 2023 | Denmark | Polygons | 4680 | 0.2m | CC-BY 4.0 | Image-annotated urban and agricultural landscapes |
| Šrollerů et al. 2025 | Czechia | Boxes | 32 | 0.02m | CC-BY 4.0 | Image-annotated high resolution data from tree-line drone imagery |
| Allen et al. 2026 | Finland, Spain | Polygons | 1609 | 0.02m | CC-BY-NC 4.0 | Fully annotated terrestrial field plots created from TLS scans |
| Hastings et al. 2020 | United States | Polygons | 1333 | 0.1m | CC-BY 4.0 | Field-verified crowns within existing terrestrial plots |

\*estimated gsd in cases where the GT data did not have spatial information.

Field-collected data are a diverse group of sources that combine information from field-based observations with image-annotated data to verify the location and extent of tree crowns. Few datasets are exclusively collected in the field due to the amount of labor needed to create datasets large enough to train machine learning models. Instead, existing tree inventories (Beery et al., 2022), and forestry surveys are often overlaid on airborne data to combine information from ground-based and airborne perspectives. Field verification may occur after initial image-annotation as in the Bornean dataset from Ball et al. 2023, and Panamanian dataset from Vasquez et al., 2023 in order to confirm existing annotations, or it may occur in parallel with multiple field-based surveys and co-registered airborne sensors as in Japanese dataset from Takeshige et al. 2025. In some cases, the airborne data is taken directly into the field on tablets to cross verify imagery with ground conditions (Graves et al., 2018). For example, Cloutier et al., 2023 delineated over 20,000 trees from Canada by stem point, which were verified using multiple drones flights and subsequent field surveys with imagery in-hand. The challenge of field collected data is associating the positions captured on the ground with the position of airborne remote sensing pixels incorporating multiple sources of error including both sensor-based uncertainty, such as geospatial location and orthomosaic generation, as well as tree biology, where tree stems often are not directly centered under tree crowns (Figure 8).

**Figure 8.**
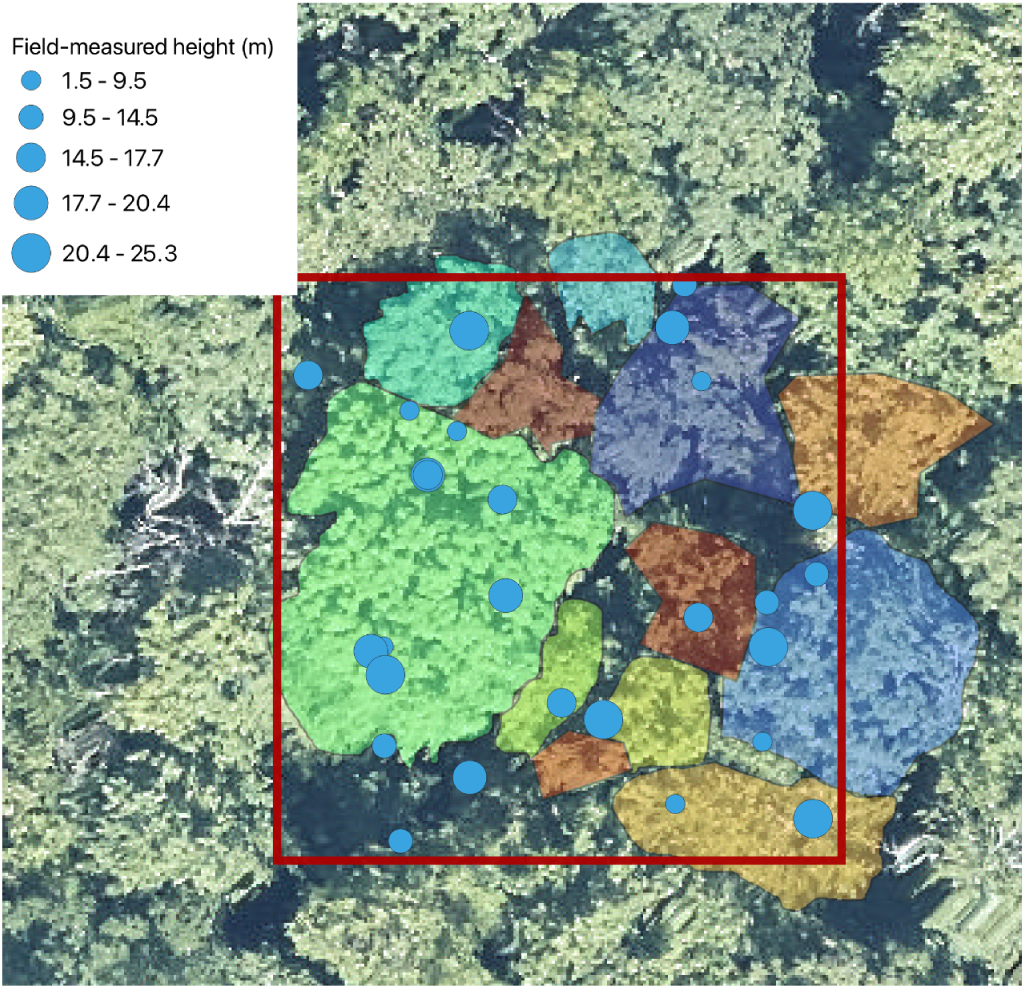
An example of the challenge of connecting field and airborne data. Field-measured points from NEON site at Harvard Forest. All trees within the red area with DBH > 10cm are tagged and measured. These data contained species, height and corresponding LiDAR derived canopy height at stem location. To create the crown delineations, the reviewer matches the location of the tree crown within the overhead imagery. Since there are many more field sampled trees than crown polygons due to overstory canopy occlusion, it is difficult to know which field-sampled points go with which overhead pixels.

In addition to data derived from human annotation and/or ground-based field data, we have also included large sources of derived annotations from simulations and unsupervised classifications. The SPREAD dataset used Unreal Engine to render realistic forest scenes and generate views from airborne perspectives. Simulated data is commonly used in other areas of remote sensing, but it has only been recently introduced to airborne tree monitoring (Rengarajan and Schott 2017). The degree to which tree detection from simulated scenes can generalize to real scenes is unknown, but holds promise for the scale of data generation and the ability to retain near perfect ground truth within the simulated scene. Similarly, we include a larger number of pseudo-labeled annotations from unsupervised models that were not directly trained for the task of RGB tree detection. The promise of unsupervised algorithm-generated annotations is the scale and diversity of images that can be accessed, thereby improving model features in a diversity of situations. The challenge is in reducing the bias generated from the upstream algorithm; this tradeoff is common in many student-teacher architectures throughout computer vision (e.g. Wang et al. 2023). The first source of weak annotations combines airborne LiDAR to extract approximate tree locations for millions of trees within the National Ecological Observatory Network. Building on Weinstein et al. (2019, 2020), LIDAR data at 27 NEON sites were processed using an existing algorithm (Silva et al., 2016) to produce approximate polygons of tree crowns. These polygons were then converted to bounding boxes based on their minimum and maximum coordinates and associated with the corresponding NEON RGB imagery (Weinstein et al., 2019, Marconi et al. 2021) and based on the needs of those projects only the bounding boxes were stored. To address issues with imperfect alignments between the data products we have now added the use of semantic segmentation models for trees (Veitch-Michaelis, 2024) to automatically identify regions where trees are present. This benchmark excludes annotations that have more than 50% non-tree pixels based on the confidence score from the semantic segmentation model. Although weak labels are inherently noisy, many studies have found that pretraining on millions of annotations yields models which perform better when fine-tuned on expert-labelled annotations compared to off-the-shelf models pretrained on general datasets or expert annotations alone (Weinstein et al., 2019, Gourde et al. 2026, Mun et al. 2026). The second source of weak labels consists of treetop points detected using canopy height models derived from photogrammetric processing of RGB drone imagery in the Open Forest Observatory (OFO) data catalog (Open Forest Observatory 2026). For all RGB imagery acquisitions (“missions”) in the OFO catalog that used a nadir camera, raw images were processed into a geospatial canopy height model and orthomosaic using a photogrammetry workflow based on Young et al. 2022. Individual canopy trees were detected as local maxima in the canopy height model using a local maximum filter implemented in the R package lidR (Roussel et al. 2020).

There is a direct tradeoff between annotation completeness on the image-level, the quality of individual tree annotations, and the diversity of out-of-sample conditions for model evaluation. Image-annotation, where trees are labeled using only one or more airborne sensors, allows for rapid complete annotation of diverse forest types, but can result in multiple trees being lumped into a single tree or a single tree being split into multiple trees (Graves et al. 2018, Allen et al. 2025). Field-based crown delineation, where either a sensor or observer is present on the ground, can produce more accurate delineations, but is time consuming and expensive resulting in fewer, often incomplete, less diverse labels. Field-stem maps, which leverage existing stem locations, can allow for rapid annotation that includes field knowledge, but can be difficult to align with airborne imagery (Graves et al. 2018) (Figure 8).

### Evaluation

We use a panel of metrics to explore the varied uses of airborne tree remote sensing represented in the benchmark data. Following Gominski et al. 2023 we adopt the terminology ‘tree counting’, ‘tree detection’ and ‘tree segmentation’. The metrics range from traditional computer vision metrics such as Average Precision, to operator metrics such as recall and precision at user-chosen score threshold. These choices reflect our goal in making metrics interpretable to applied projects that need to choose a specific cutoff for tree prediction to be used in downstream analysis.

#### Visible Tree Counting

Tree counting is a common task of airborne remote sensing that estimates the number of individual trees in a given area visible from above (i.e., in the canopy including small sun-exposed trees in canopy gaps). For counts, we use the mean absolute error between the ground-truth count and the predicted count per image as our evaluation metric.

#### Tree Detection

Tree detection is the task of locating the individual tree within an image. It is usually accompanied with a measure of goodness of fit by overlaying area-based predictions, such as the intersection-over-union of boxes or polygons over ground-truth annotations. Given a model specific confidence threshold (min_score), series of ground truth boxes per image (G), a set of predictions per image (P), a minimum prediction confidence score, and a degree of overlap, we calculate a tree detection recall and precision.

Step 1: Filter predictions by confidence

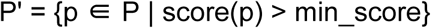

Step 2: Calculate IoU for all prediction-ground truth pairs

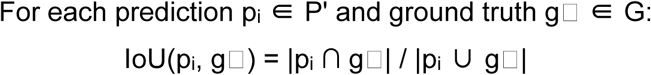

Step 3: Determine matches

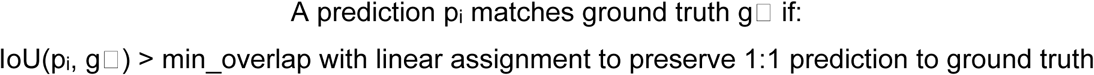

Step 4: Calculate recall

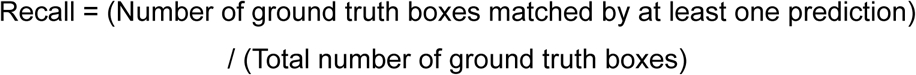

When the ground truth image is fully annotated we also calculate precision:

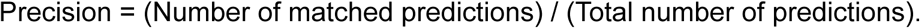

Step 5. Combining these metrics into a single F1 metric

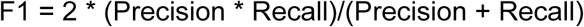

In the case of TreePoints, we cannot measure overlap, since points have no area. Instead we use a radius around each point based on a fixed euclidean distance of 4m to be considered a true positive match. A predicted tree center can only match to one true positive ground truth location using Hungarian matching.

For incompletely annotated datasets, it is not possible to differentiate low precision predictions from true-positive examples that have not been annotated. Instead of calculating only recall for incompletely annotated evaluation datasets, we calculate Mask-aware Precision. Mask-aware precision aids in calculating precision in cases of partially labeled imagery by not penalizing for predictions made in areas of tree pixels but lacking ground truth (Figure 9). Using the tree versus background mask from Veitch-Michaelis et al. 2024, we use the predicted class of each pixel within a tree prediction to only penalize those predictions that do not match a ground truth and have less than 50% tree pixels. This prevents us from penalizing a true positive prediction that is missing the associated ground truth. While this model-assisted approach is imperfect, it is superior to solely calculating recall, since an algorithm that produced a large number of predictions per image is likely to have at least some predictions that overlap with ground truth. In the rare cases that the tree versus background mask disagreed with ground truth, we manually reviewed the associated images and either removed or maintained the annotations based on visual interpretation.

**Figure 9.**
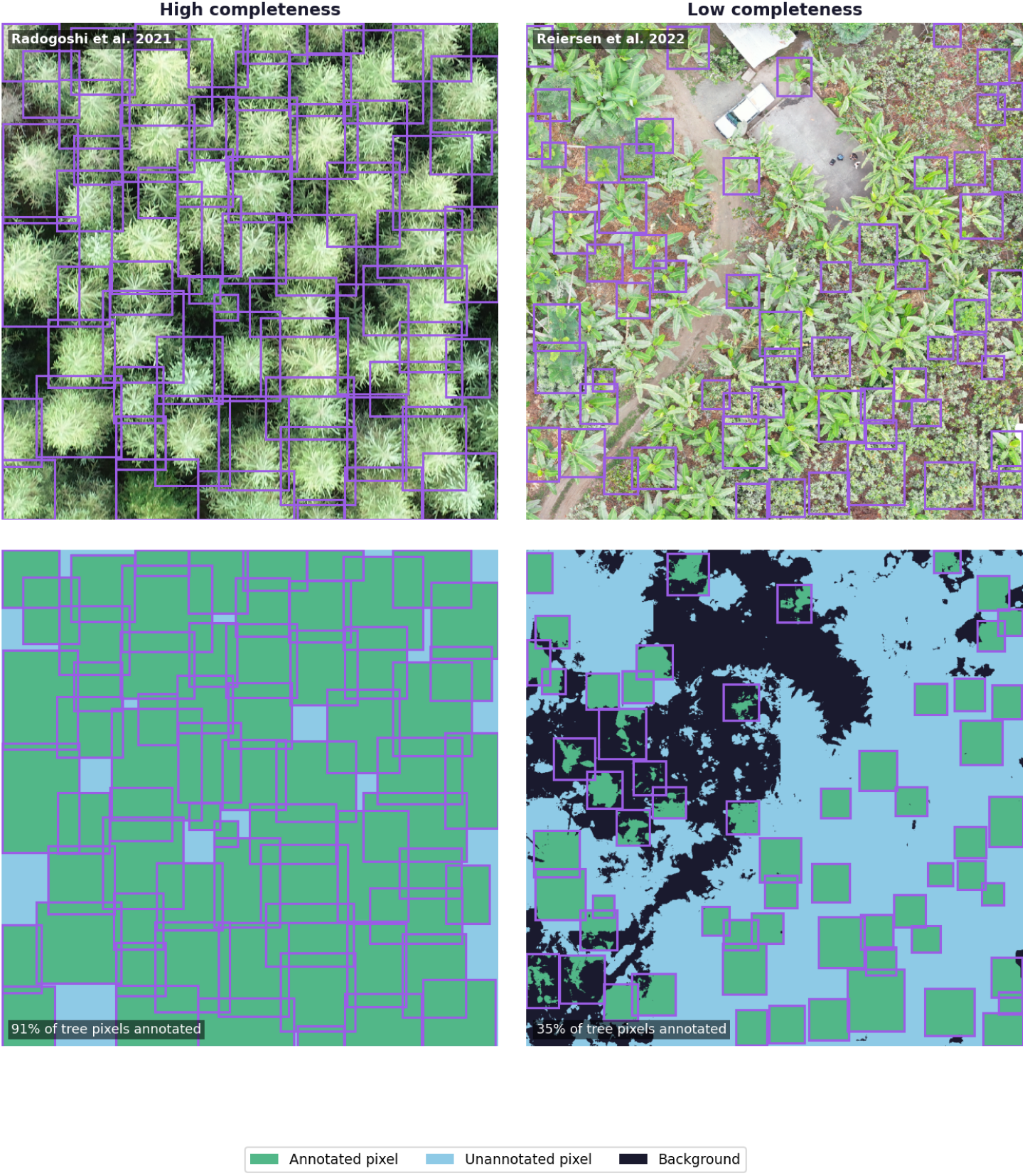
Each image in the dataset is accompanied by an annotation mask layer that uses a semantic segmentation model (Veitch-Michaelis et al. 2024) to predict tree versus background pixels. From this layer we can differentiate annotated versus unannotated portions of tree cover, which is a key limitation of many datasets.

We calculate the F1 score (harmonic mean of precision and recall) with an IoU threshold of 0.4 as our primary metric of model performance. We focus on F1 with a fixed threshold because most applications of computer vision involving trees require a clear designation of whether an object is a tree or not and this threshold has been the most effective choice in previous work (Weinstein et al. 2019, 2020). In addition to F1 we also calculate Average Precision (AP), a standard metric in computer vision that combines precision and recall at different score thresholds by integrating over precision-recall curve to calculate the area under the curve across thresholds as a measure of confidence weighted accuracy. It is similar to a receiver operator curve (ROC) and is a single measure that captures both the true positive rate, the false positive rate, and the confidence scores. To provide a more refined measure of crown boundaries, we calculate AP@60 for the validation dataset as well as AP@40. The choice of these IoU levels is arbitrary and relates to the varied use cases for tree detection predictions. The AP@60 measure is strict enough that it likely surpasses the level of granularity possible through human annotation of airborne imagery and is most appropriate for the validation TLS datasets. Visual review of IoU and the validation data suggests that true positives at the 0.4 level are generally correct, but lack fine boundaries, whereas 0.6 are highly aligned between ground truth and prediction and appropriate for more fine-grained tasks that require strict crown area agreement (Figure S1).

Tree segmentation aims to define the boundaries of individual tree crowns visible in the airborne imagery. We use the same metrics as tree detection for evaluating segmentation. Tree segmentation is the most common form of evaluation for traditional computer vision approaches, since it connects to well-established approaches in the wider machine learning literature. However, it should be noted that the airborne perspective of individual tree segmentation complicates the fine-scale annotation of object boundaries, because crowns can be interdigitated and it requires defining the edge of a three dimensional tree crown in a two dimensional image view.

#### Benchmarking Existing Models

We provide baseline models to set the minimal performance for each task using standard off-the-shelf computer vision tools and existing tree detection algorithms. Our goal here is not to set a new standard of performance, but to codify the current state of existing models to better organize and clarify the next steps for global tree detection. We limited our exploration of models to existing published approaches and we briefly describe the models here and provide citations for each for those interested in additional detail.

##### DeepForest NEON Tree Detector

The DeepForest Tree Box Detector is an object-detection model that uses a RetinaNet architecture to predict canopy tree bounding boxes. DeepForest was trained using a combination of large-scale LiDAR-derived pretraining annotations and image-annotated crowns, predominantly from forested habitats in the National Ecological Observatory Network. For more information on model training, generalization, and application see Weinstein et al. 2019, Weinstein et al. 2020.

##### SAM2 and SAM3

Segment-anything is a foundation computer vision model developed by Meta that uses promptable transformers to segment image pixels into classes. While they are similarly named, SAM2 focused on using geometric prompts to create individual instance segmentations, and SAM3 introduced text-based semantic prompts for open set segmentation. Both models use extremely large pretrained transformers to map custom natural language inputs, such as ‘individual tree canopy’ into segmentation masks and individual object boundaries. SAM3 is trained on a large corpus of data, and it is unknown whether any tree detection tasks exist in their data pipeline. SAM2 and SAM3 are respectively released by Meta under the Apache 2.0 license and a custom SAM license for commercial and scientific applications.

##### Mask R-CNN

Mask R-CNN is a common instance segmentation algorithm that uses pixel-level masks to generate polygon predictions based on convolutional image features (He et al. 2018). Mask R-CNN is easy to train and serves as a reasonable baseline for polygon training and prediction using the flagship torchvision Python package. While the performance may be lower than transformer-based architectures (e.g. Schmid et al. 2025), Mask-RCNN has been commonly used for polygon inference in tree detection (e.g. Ball et al. 2023) and serves as off-the-shelf baseline for MillionTrees polygon tasks. To compare with DetecTree2, we used the Detectron implementation of Mask R-CNN.

##### CenterNet

CenterNet takes a different approach to object detection by reformulating the task as keypoint estimation, identifying object centers directly in a heatmap rather than relying on region proposals or anchors (Zhou et al. 2019). The simplicity of this center-point representation makes CenterNet computationally efficient and straightforward to implement in native Pytorch with the ubiquitous UNet-ResNet50 architecture. Though newer transformer-based detectors have achieved superior performance on benchmark datasets, CenterNet remains practical for tree detection workflows—particularly where computational efficiency or ease of deployment is prioritized—and has seen adoption in remote sensing applications (Gominski et al. 2026, Brandt et al. 2024).

##### CanopyRS

CanopyRS is a Python library providing pretrained, general-purpose deep learning models and inference-ready pipelines to detect and segment trees in remote-sensing imagery. First introduced alongside the SelvaBox tropical dataset (Baudchon et al., 2026), it provides a DINO detection transformer (Zhang et al., 2023) trained on a mix of four datasets spanning tropical, temperate and urban biomes with multi-resolution augmentations. It was later extended with a SAM3 checkpoint finetuned on SelvaMask (Duguay et al., 2026), a densely annotated tropical tree crown segmentation dataset. Several of CanopyRS’s training datasets are also included in MillionTrees, particular care was taken to ensure no overlap between CanopyRS’s training data and the MillionTrees test splits. For this reason, the CanopyRS model was not assessed for the out-of-distribution scores since a portion of the sources used for these tests were already present in these existing model weights. For the box geometry, we use the DINO detector specified in the ‘preset_det_multi_NQOS_dino_swinL’ config for the tree detection task, and for polygon geometry we pair a DINO detector and SAM3, both finetuned on SelvaMask, taking their model definitions from the config ‘preset_seg_multi_NQOS_selvamask_SAM3_FT_quality’. All models and configs can be found in the CanopyRS huggingface repository.

##### TreeFormer

TreeFormer is a point transformer-based architecture that uses a combination of losses to optimize the number of tree predictions within the entire image with the local features of predictions using prediction density maps (Amirkolaee et al. 2023). For the existing model comparison we implemented the TreeFormer architecture and pretrained it on the NEON weakly labeled data included in the MillionTrees box dataset. Boxes were transformed into points using the centroid. The architecture was re-implemented in the DeepForest Python package along with the hyperparameter information to reproduce our model training regime. To act as an ‘existing’ model baseline for the TreeFormer point detector, we trained the architecture on the ‘KCL London’ dataset included in MillionTrees that was connected with the original TreeFormer manuscript.

##### DetectTree2

DetectTree2 is a tree crown polygon prediction model based on Meta’s Detectron Mask R-CNN implementation (Ball et al. 2023). It was trained using data from Borneo and French Guiana, with a portion of that data present in MillionTrees training sources. DetectTree2 used 3797 crowns to train models based on a combination of image labeling and field verification and assessed both within domain scores as well as multiple temporal views to assess generalization through time.

## Data Availability

The data in this benchmark is available through the MillionTrees python package. The source code for the python package is open source on github (https://github.com/weecology/MillionTrees). A static version of the release dataset at time of submission is submitted on Zenodo.

## Acknowledgments

The creation of the MillionTrees benchmark was supported by the World Resources Institute and its donors the Bezos Earth Fund and The Audacious Project. This work was also supported by the USDA National Institute of Food and Agriculture through the Hatch project FLA-WEC-006662. and McIntire Stennis project FLA-FOR-006695. M. J. Allen and S. H. M. Hickman was supported by the UKRI Centre for Doctoral Training in Application of Artificial Intelligence to the study of Environmental Risks [EP/S022961/1]. E. R. Lines, S. W. D. Grieve and H. J. F. Owen was funded by a UKRI Future Leaders Fellowship awarded to E. R. Lines [MR/T019832/1]. T. Kattenborn acknowledges support via Germany’s Excellence Strategy (DFG Future Forests–EXC-3127-533786343). Contributions by D. J. N. Young and D. J. Russell were supported by the U.S. National Science Foundation [#2152671]. S. Beery was supported by an NSF CAREER Award [#2441060]. M. Mugabowindekwe acknowledges funding from the European Research Council (ERC) under the European Union’s Horizon 2020 Research and Innovation Programme [grant agreement no. 101157661, TREEMAP] and from the Danish National Research Foundation Center for Remote Sensing and Deep Learning of Global Tree Resources (TreeSense) [DNRF192]. B. Wagner was supported by the Australian Research Council [LP140100580, FT120100715] and by the University of Melbourne through a Melbourne Research Scholarship, with additional funding from the Royal Zoological Society of New South Wales, the Foundation for Parks and Wildlife, the Norman Wettenhall Foundation, the former School of Ecosystem and Forest Sciences at the University of Melbourne, and the Integrated Forest Ecosystem Research (IFER) program of the Victorian Department of Energy, Environment and Climate Action (DEECA). Y. Sun was supported by the National Natural Science Foundation of China [42571405]. Y. Onoda acknowledges support via JSPS KAKENHI [21H05314, 21H02564]. J. Troles was supported by the Bavarian Ministry of Digital Affairs and the Federal Ministry for Housing, Urban Development and Building.

---

**Figure S1.**
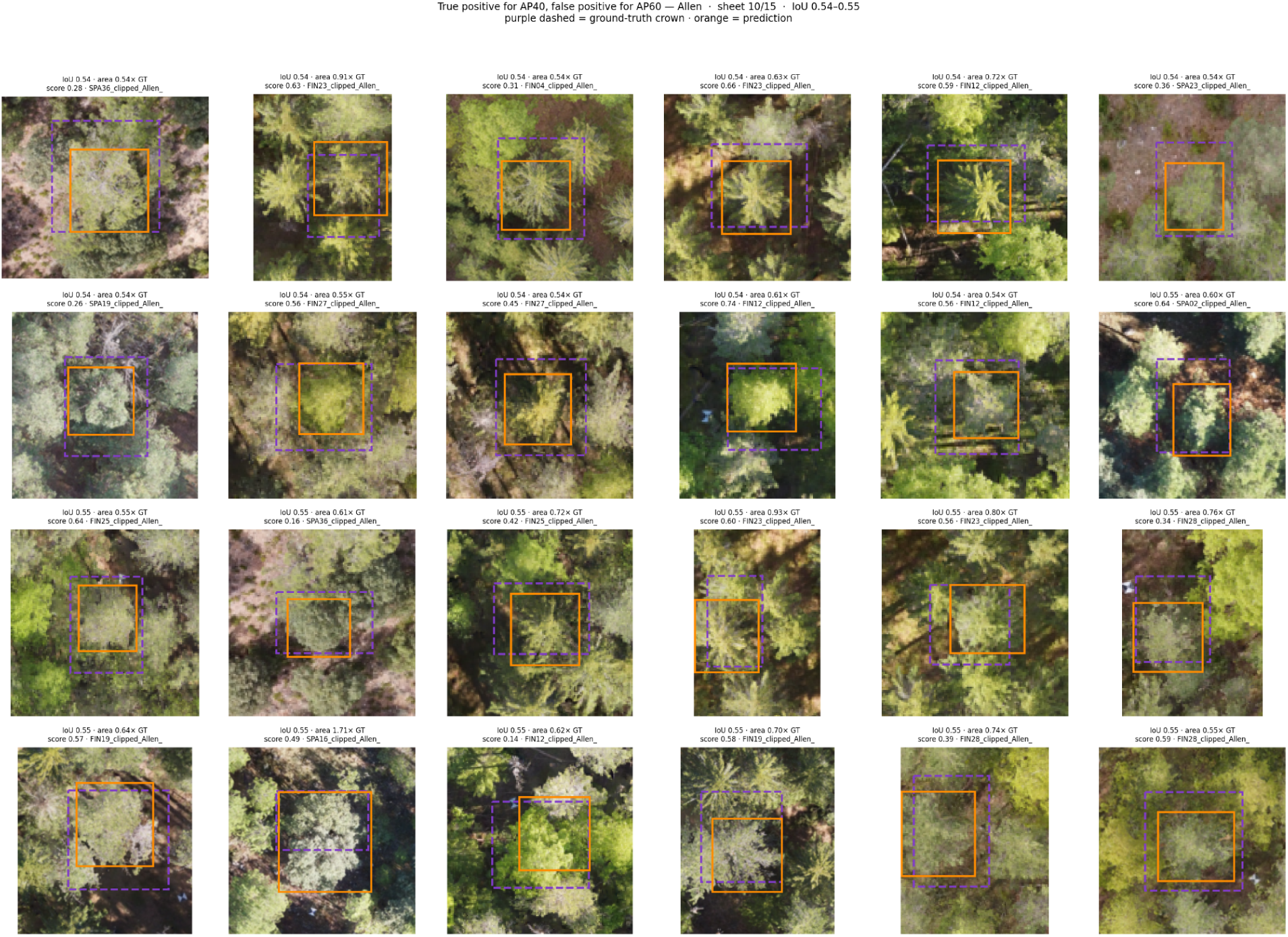
Visual inspection of intersection-over-union between ground truth and the predictions from the fine-tuned DeepForest model. Predictions were sorted by IoU, and predictions with IoU between 0.5 and 0.6 are shown below.

